# AP-1 activation in *Drosophila* neuropil ensheathing glia improves traumatic brain injury survival

**DOI:** 10.64898/2026.08.13.744727

**Authors:** Michael Fetchko, Shambhavi Gupta, Seanna E. Kelly, Akanksha S. Mathivanan, Stephen W. Ratner, Shorbon Mowla, Namarata Battula, Maryem H. Abdelgelil, Annika F. Barber

**Affiliations:** Waksman Institute, Rutgers University, 190 Frelinghuysen Rd., Piscataway, New Jersey, 08854; School of Arts and Sciences, Rutgers University, 77 Hamilton Street, New Brunswick, NJ 08901; Department of Molecular Biology and Biochemistry, Rutgers University, 604 Allison Road, Piscataway, NJ 08854

**Keywords:** *Drosophila*, glia, traumatic brain injury, blood-brain barrier, AP-1

## Abstract

Traumatic brain injury (TBI) impacts millions of individuals annually causing death, disability, and a heightened risk for long-term neurological and neuropsychiatric disorders. In recent years the fruit fly, *Drosophila melanogaster* has become a valuable model organism to study the cellular and molecular responses following TBI. AP-1 mediated transcriptional responses to TBI have previously been identified in *Drosophila* using pan-glial approaches. Fruit flies possess multiple glial subtypes which vary greatly in both cellular morphology and function, including glia of the blood hemolymph barrier, cortex, astrocyte-like, and ensheathing glia. By generating and utilizing a nuclear localized AP-1 transcriptional reporter, we identified glial subtype-specific differences in the extent of AP-1 activation following injury. Our findings identify a strong AP-1 response in the blood hemolymph barrier and ensheathing glia, a moderate response in cortex glia and little to no AP-1 activation in astrocyte-like glia. In addition, we inhibited AP-1 signaling in each glial subtype and tested the effect on acute survival. We found that inhibition of the AP-1 response in neuropil ensheathing glia leads to increased mortality following mild and moderate TBI. These results show that AP-1 activation levels vary across glial subtypes after TBI, with activation in neuropil ensheathing glia having a particularly important role in promoting post-injury survival.

**ARTICLE SUMMARY:** Using *Drosophila* as a model organism, we investigated the early molecular and cellular response to traumatic brain injury. Our findings substantiate the requirement of a functional glial associated AP-1 transcriptional activation response for survival. Using colocalization studies, we characterized the AP-1 glial response in six morphologically and functionally distinct glia subtypes. After TBI, we find high levels of AP-1 activation in glia of the hemolymph brain barrier, cortex glia, and ensheathing glia. We further show the importance of AP-1 transcription within the neuropil ensheathing glia subtype for optimal survival following TBI.

## INTRODUCTION

Annually, estimates of 2.8 million new cases of traumatic brain injury (TBI) occur and currently over 5 million individuals suffer chronic brain injury-related disabilities in the United States (Faul et al. 2010; Peterson et al. 2019). Lasting effects of brain injury include chronic pain, depression, impaired memory, increased dementia risk, and a shortened lifespan of on average 7-9 years (Kumari et al. 2025). While improved treatments have resulted in decreased mortality after TBI, there have been no major breakthroughs that address the chronic impairments experienced by survivors. The pathology of TBI is highly complex, which has limited development of clinical approaches to diagnose, predict, and treat post-injury pathologies. Due to the limited understanding of the underlying cellular and molecular causes of post-TBI sequelae, many clinical interventions focus on symptom relief. A better understanding of the cellular and molecular response to head injury is necessary to develop precision interventions that improve quality of life for TBI survivors.

Animal models have greatly refined our understanding of how organisms respond to and recover from head injury at the cellular and molecular levels (Krishnamurthy and Laskowitz 2016; Kumari et al. 2025). However, pharmacological approaches that have shown promise in rodent models have thus far failed in human clinical trials. The fruit fly, *Drosophila melanogaster*, has been a workhorse of molecular genetics for decades, with high conservation of genes, molecular pathways, and cell types from *Drosophila* to humans enabling translatability of many findings (Yamamoto et al. 2014). There are multiple models of fly TBI, which have contributed extensively toward uncovering genes and signaling pathways involved in the TBI response, including genetic factors also identified in human patients (Katzenberger et al. 2013; Anderson et al. 2021; Byrns et al. 2021; Aggarwal et al. 2022; Kelly et al. 2025). Fly TBI paradigms may involve whole-animal or head-specific trauma with varying levels of throughput and calibration (Aggarwal et al. 2022; Kelly et al. 2025). In both flies and mammals, an activator protein 1 (AP-1) mediated transcriptional response is upregulated and associated with glial cells after head-specific TBI. In humans, AP-1 transcriptional activity is detected after moderate TBI and correlates with phagocytic microglial activation (Byrns et al. 2021). Similarly, RNAseq analysis revealed that flies activate a massive glial AP-1 transcriptional response which is essential for survival after TBI (Byrns et al. 2021).

The AP-1 transcription factor family of basic leucine zipper (bZIP) transcription factors namely *fos*, *jun* and the activating transcription factor (ATF) family, are highly homologous, evolutionarily conserved factors which regulate cellular processes including proliferation, differentiation, and apoptosis (Hess et al. 2004). Activation of AP-1 is largely regulated by Mitogen-Activated Protein Kinase (MAPK) pathways which are triggered by secreted cytokine and growth factor proteins in response to immune stimuli, stress, or during normal development. Active MAPK phosphorylates individual bZIP subunits which enables formation of bZIP homo- or heterodimers that bind DNA regulatory elements, called TPA-responsive elements (*TRE*s), and modulate gene transcription (Angel et al. 1987; Lee et al. 1987). Transgenic flies containing the fluorescent AP-1 reporter *TRE*_*dsRed* upregulate reporter expression in glial cells following TBI (Chatterjee and Bohmann 2012; Byrns et al. 2021).

In *Drosophila*, glial cells are essential nervous system support cells which have been identified in tissues by their expression of the pan-glial homeodomain transcription factor gene, *reversed polarity* (*repo*) (Xiong et al. 1994; Kremer et al. 2017). *Drosophila* glia can be divided into five major subtypes: perineurial glia (PG), subperineurial glia (SPG), cortex glia (CG), astrocyte-like glia (ALG), and ensheathing glia (EG), which differ considerably regarding cell size, shape, function and location within the nervous system (Zwarts et al. 2015; Kremer et al. 2017). PG and SPG constitute the *Drosophila* hemolymph-brain barrier (HBB) which serves the same function as the mammalian blood brain barrier; however, insects use hemolymph whereas mammals utilize blood as liquid transport for nutrients, waste and cellular signals (Stork et al. 2008; Dunton et al. 2021). PG form the outermost surface glia layer and generate a semipermeable protective membrane which regulates nutrient and metabolite flow and directly contacts the hemolymph (Leiserson et al. 2000). Below the PG, SPG utilize tight septate junctions to arrange in a single cell layer which inhibits the paracellular flow of large potentially harmful macromolecules into the nervous system. CG are the only glial subtype found within the cortical region of the brain located just beneath the SPG layer of the HBB. Most neuron cell bodies and nuclei are also located within the cortex where single cortex glia send out flat cellular projections called lamellae which envelop multiple neuron cell bodies. CG are known to physically compartmentalize neurons as well as provide trophic and neuromodulatory factors for neuron function (Bittern et al. 2021).

*Drosophila* ALG are neuropil associated glia where individual ALG cover the neuropil surface in a non-overlapping fashion. From the surface of the neuropil, ALG send projections deep into the neuropile to interact with neurons in the synaptic spaces (Stork et al. 2014). ALG are known to be involved in synaptic clearance, developmental sculpting and metabolic support. Ensheathing glia, like ALG, are neuropil-associated glia (Kremer et al. 2017; Pogodalla et al. 2022). EG wrap around axon bundles, dendrites and entire brain structures such as olfactory bulbs and the ellipsoid body. EG serve to insulate neuronal circuits from one another and also as the brain’s primary debris clearance mechanism of apoptotic or injured cells (Doherty et al. 2009; Bittern et al. 2021).

Previous studies in *Drosophila* have broadly characterized pan-glial AP-1 response after TBI (Byrns et al. 2021). Using fluorescent nuclear cell markers, our work establishes that glial subtypes differentially activate AP-1 24 h following TBI and that AP-1 activation occurs strongly in the HBB glia and EG, with a moderate response in CG and negligible AP-1 activation in ALG. Furthermore, we show that inhibition of AP-1 signaling in all glia and specifically in neuropil ensheathing glia (NEG) increases mortality following TBI. We find that inactivating AP-1 in NEG reduces the number of NEG per brain which may underlie the mortality increase observed after injury. These results suggest different transcriptional outputs of the AP-1 pathway depending upon its cellular context and implies AP-1 activation coordination amongst glia subtypes is required following brain injury for optimal survival.

## MATERIALS AND METHODS

### Fly stocks and husbandry

See the reagent table (Table S1) for the list of fly stocks used in this manuscript. Flies were maintained on a standard Bloomington *Drosophila* Stock Center cornmeal diet and cultured at 25° C with 50-70% humidity.

#### TBI Administration

Injuries were performed as previously described in (Saikumar et al. 2021; Ratner et al. 2024). Prior to injury, male flies were collected at 1-2 days post-eclosion and housed 18 flies per vial in a 12:12 light-dark cycle at 25° C for 3 days. All injuries were performed between ZT 0-4. To injure, 4-5 day old flies were briefly anesthetized with CO_2_ and loaded 6 flies per collar. Collars were placed underneath the piezo until the edge of the piezo aligns where the proboscis meets the head (Figure 1a). The device voltage is then set to the desired injury severity, and a single crush injury is administered to each collar. After injury, collars are slid from beneath the piezo and flies are unloaded from the collar by sliding a forceps along the underside groove of the collar. Flies are then placed into horizontal food vials for a 1–2-hour recovery period. For brain imaging experiments, uncollared shams were used to eliminate any signal due to rough handling. For survival assays, sham flies were placed in collars for approximately 2-3 minutes and then removed with forceps, as described above.

**Figure 1.**
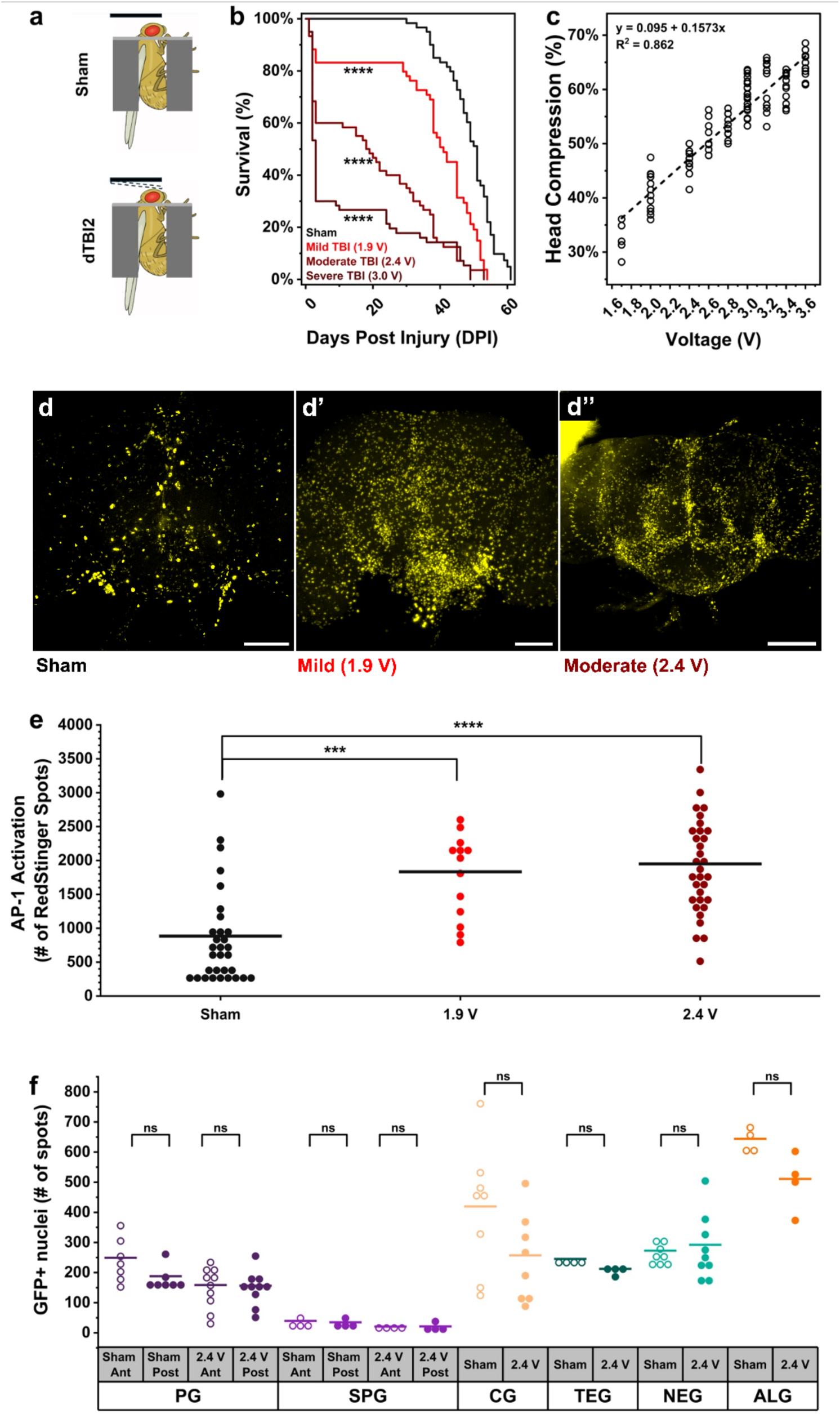
dTBI2 drives severity-dependent effects on fly survival and AP-1 pathway activation. a) Schematic of dTBI2 device sham (top) and injury (bottom) modalities. b) Kaplan-Meier survival curve of full lifespan assay for sham (black), mild 1.9 V injury (light red), moderate 2.4 V injury (red), and severe 3.0 V injury (dark red) TBI flies. Log-rank pairwise comparison was used to determine statistical differences between sham controls and each injury group. N = 60 flies per condition, one biological replicate. c) Quantification of percent head compression at increasing device voltage with linear regression analysis, R^2^ = 0.86. N = 1-3 collars per voltage, 6 flies per collar, each circle represents an individual fly. d-d’’) Representative images of *TRE*_*RedStinger* activation (yellow) in the central brain for sham (d), mild (d’), and moderately (d’’) injured flies. e) Quantification of total number of *TRE*_*RedStinger* spots in the brains of sham, mild, and moderately injured flies shows a significant increase of AP-1 activation with increasing injury severity. Each dot represents one brain. f) Quantification of GFP positive nuclei within the brains of each glial subtype before and after injury shows no change in the number of glial cells after injury. Each dot represents one brain for the indicated subtypes with sham noted with open circles and injured flies noted with closed circles. N = 4-10 brains per subtype and condition. Significance was assessed by 1-way ANOVA with a Tukey’s *post hoc* test was used to determine statistical significance between sham and TBI conditions. *** p<0.001, **** p<0.0001, ns = not significant

#### Survival Assay

All genotypes used in survival assays were backcrossed to our isogenic wild-type control line, *Iso*^31^, at least 5 times. Following injuries, flies were housed 18 per vial, mortality was counted daily, denoting how many flies per vial were alive, dropped, and dead. Throughout the duration of the assay, survivors were flipped onto fresh food every second day. For full lifespan experiments, the assay continued until all control flies died. For survival assays, sham flies were loaded into the collar and subsequently unloaded without injuring using the piezo. Survival assay data was analyzed using Kaplan-Meier analysis with log-rank comparison in OriginLab 2025b.

#### Smurf Assay

A dye-feeding assay was used to assess gut permeability based on the original assay (Wong et al. 2009) and those used in other TBI paradigms (Katzenberger et al. 2015; Barekat et al. 2016). To increase male dye intake, we used a less nutritive diet that required flies to eat larger volumes of food, and hence dye, to meet caloric needs. Less nutritive food consisted of 12.5 g of corn meal, 1.5 g of agar, 0.75 g of yeast, and 138 mL of water, dyed with 1.875 g (2.5%) of erioglaucine disodium salt. Male and female flies were collected at 3 days post-eclosion and placed on undyed or dyed less nutritive food 48 h prior to, sham, moderate (2.4 V) dTBI2, or mixer mill TBI model (Barekat et al. 2016) for 30 seconds at 100 rpm and placed back on their original food source. Twenty-four hours post-injury all flies were imaged using a Zeiss microscope camera. Gut barrier integrity was determined by observing whether the dye stayed consolidated in the gut (not smurfed) or the dye spread throughout the abdomen and body (smurfed).

### Generating pH_TRE_RedStinger_attB

*pH_TRE_RedStinger_attB* was designed to be identical to the *P{TRE-DsRedT4}* (Flybase ID-FBtp0072198) construct within the effective reporter region of the constructs. Differences occur within the plasmid backbones outside of the *gypsy* transcriptional insulators that flank the reporter region (Chatterjee and Bohmann 2012). *pH_TRE_RedStinger_attB* was generated in a two-step process by first making the *pH_RedStinger_attB* plasmid containing a *ϕ*C31 *attB* site, the nuclear localized *RedStinger* open reading frame, a *miniwhite* gene, and an *origin/AmpR*, but lacking the *TRE* repeats. *ph_RedStinger_attB* was assembled through a 5 PCR fragment NEBuilder® reaction. See Table S2, for the primers and templates used for PCR.

To add the *TRE* sequence to *pH_RedStinger_attB*, *TRE* sequence was amplified from genomic DNA of *TRE*-*DsRed* fly genomic DNA using the following primer combination: TRE_seq_F2 and TRE_R_XhoI (Table S2). An NEBuilder® reaction was used to insert the amplified *TRE* containing fragment into the *pH_Red_Stinger_attB* plasmid that was cut with *SphI* and *AgeI.* The resulting *pH_TRE_RedStinger_attB* plasmid was integrated into flies at the *attP40* landing site using *ϕ*C31 integrase (BestGene Inc. CA, USA).

#### Brain dissection and confocal imaging

A brain dissection method to preserve the brain hemolymph barrier was adapted from (Bhasiin et al. 2023). Twenty-four hours following injury, whole flies were submerged in PBST (PBS + 0.1% Tween-20). PBST was removed and the flies were fixed for 25 minutes in 4% paraformaldehyde/PBST at room temperature (RT). Flies were washed in PBST and their proboscises were removed prior to a second 4% paraformaldehyde fix for 10 minutes at RT. After the second fix, fly brains were dissected from the cuticle, washed once with PBST, and mounted in Vectashield®.

Brains expressing RedStinger and GFP.nls were imaged using a 20x oil objective via sequential scans on a Leica TCS SP8 Lng True Confocal Point Scanning (TCS) Microscope using the 552 and 488 nm lasers, respectively. Both step size and pixel resolution were optimized using the Leica software with a section step size ranging from 1.00-1.25 μm and resolutions ranging from 4700x4700-5300x5300 pixels. Images were scanned bidirectionally with a line average of 3. Prior to analysis, images were deconvolved using Leica’s “lightning deconvolution”.

### Confocal image processing and colocalization analysis

Each brain image was processed in FIJI to form substacks in the z-plane for analysis (Schindelin et al. 2012). Unique substack regions were selected for each glial subtype to optimize the number of glia for tractable counting. See Table S3 for selected substack regions for each glial subtype. Colocalization analysis was performed in Bitplane Imaris v. 10.2.0 with an ROI drawn manually around the central brain using the built-in surface feature. We used the built-in Imaris “Spots” function to independently identify GFP^+^ (settings varied by glial subtype, see Table S4, manual thresholding) and RedStinger^+^ (x-y diameter = 5 μm, z spread = 11 μm, manual thresholding) nuclei. After spot assignment, spots were colocalized using the Spot-Spot colocalization extension with 2.5 μm as the maximum distance between spot centers. Statistical significance comparing the percentage of colocalized nuclei between sham and injured brains was determined by performing a one-way ANOVA with Tukey’s *post hoc* test. The number of glia subtype cells per sample analyzed during colocalization studies is presented in Table S5.

## RESULTS

### Severity-dependent effects of dTBI2 on *Drosophila* lifespan and brain AP-1 activation

We administered traumatic brain injury to 5-day old male isogenic *w*^1118^ flies using the dTBI2 method which uses a piezoelectric plate to deliver head compression to immobilized, unanesthetized flies (Fig. 1 a) (Saikumar et al. 2021; Ratner et al. 2024). We defined injury severity based on median lifespan after injury, with mild, moderate, and severe injury significantly reducing lifespan compared to sham control flies which are placed into the immobilization collar but not injured (Fig. 1 b). As in other TBI paradigms, mortality after injury occurs in two phases: an initial period of high mortality in the first 48 h post injury followed by a slower phase of mortality from then on. At mild injury severity, the slope of the mortality curve is similar to that of sham controls after initial high mortality, while at more severe injury levels both mortality phases are distinct from sham controls. Prior studies using other TBI administration methods have found that substantial mortality arises from reduced gut barrier integrity and consequent inflammatory responses after injury (Katzenberger et al. 2015; Barekat et al. 2016; Scharenbrock et al. 2021). Using the dTBI2 paradigm, no flies exhibited leakage of blue dye from the gut into the periphery 24 h post-injury, while 29% of flies injured using a mixer mill TBI paradigm (Barekat et al. 2016) showed the blue dye-leakage “smurf” phenotype 24 h post-injury (Fig. S1) (Rera et al. 2012). Thus, we conclude that dTBI2-induced acute mortality does not result from impaired gut barrier integrity. The amount of head compression is determined by the voltage through the piezoelectric activator, which is linear and reproducible across individual flies (Fig. 1c). Using these metrics, we defined mild injury as occurring at 1.9 V with an average head compression of 29% ± 0.4% and a median lifespan of 40 days, moderate injury as occurring at 2.4 V with an average head compression of 46% ± 0.6% and a median lifespan of 20 days, and severe injury as occurring at 3.0 V with an average head compression of 59% ± 0.6% and a median lifespan of 3 days.

Prior studies report increased activation of the AP-1 signaling pathway in fly glia after TBI using a *TRE*_*dsRed* reporter of AP-1 activation (Chatterjee and Bohmann 2012; Byrns et al. 2021). To facilitate colocalization analysis with other nuclear reporters, we created the *TRE*_*RedStinger* reporter in which *TRE*-elements (Risse et al. 1989) are fused to the nuclear-localized *RedStinger* reporter sequence (Risse et al. 1989; Barolo et al. 2004; Chatterjee and Bohmann 2012). Our reporter recapitulates prior findings of injury-severity-dependent activation of AP-1 signaling (Fig. 1 d-e) (Byrns et al. 2021). At baseline, AP-1 activation is reported by *TRE*_*RedStinger* expression in 883 ± 125 cells per brain (Fig. 1 d, e). When assayed 24 h after injury, the number of RedStinger^+^ cells increased significantly compared to sham controls with an average of 1833 ± 172 cells after mild injury and 1950 ± 120 cells after severe injury (Fig. 2 d’, d’’, e). To assess whether TBI induces large scale cell death, we drove expression of nuclear GFP reporter in six glial subpopulations using the Gal4-UAS system and compared the number of GFP^+^ nuclei at 24 h post-injury to sham controls (Fig. 1f). We assessed GFP expression in six glial subpopulations: perineurial glia (GMR85G01-GAL4, termed PG-Gal4), subperineurial glia (GMR54C07-GAL4, termed SPG-Gal4), cortex glia (GMR54H02-GAL4, termed CG-Gal4), tract ensheathing glia (GMR75H03-GAL4, termed TEG-Gal4), neuropil ensheathing glia (GMR56F03-GAL4, termed NEG-Gal4), and astrocyte-like glia (GMR86E01-GAL4, termed ALG-Gal4) (Pfeiffer et al. 2010; Kremer et al. 2017). We found no significant difference in the number of GFP-labeled glia for any glial subpopulation at 24 h post injury, suggesting that there is no substantial glial cell death at this time point.

**Figure 2.**
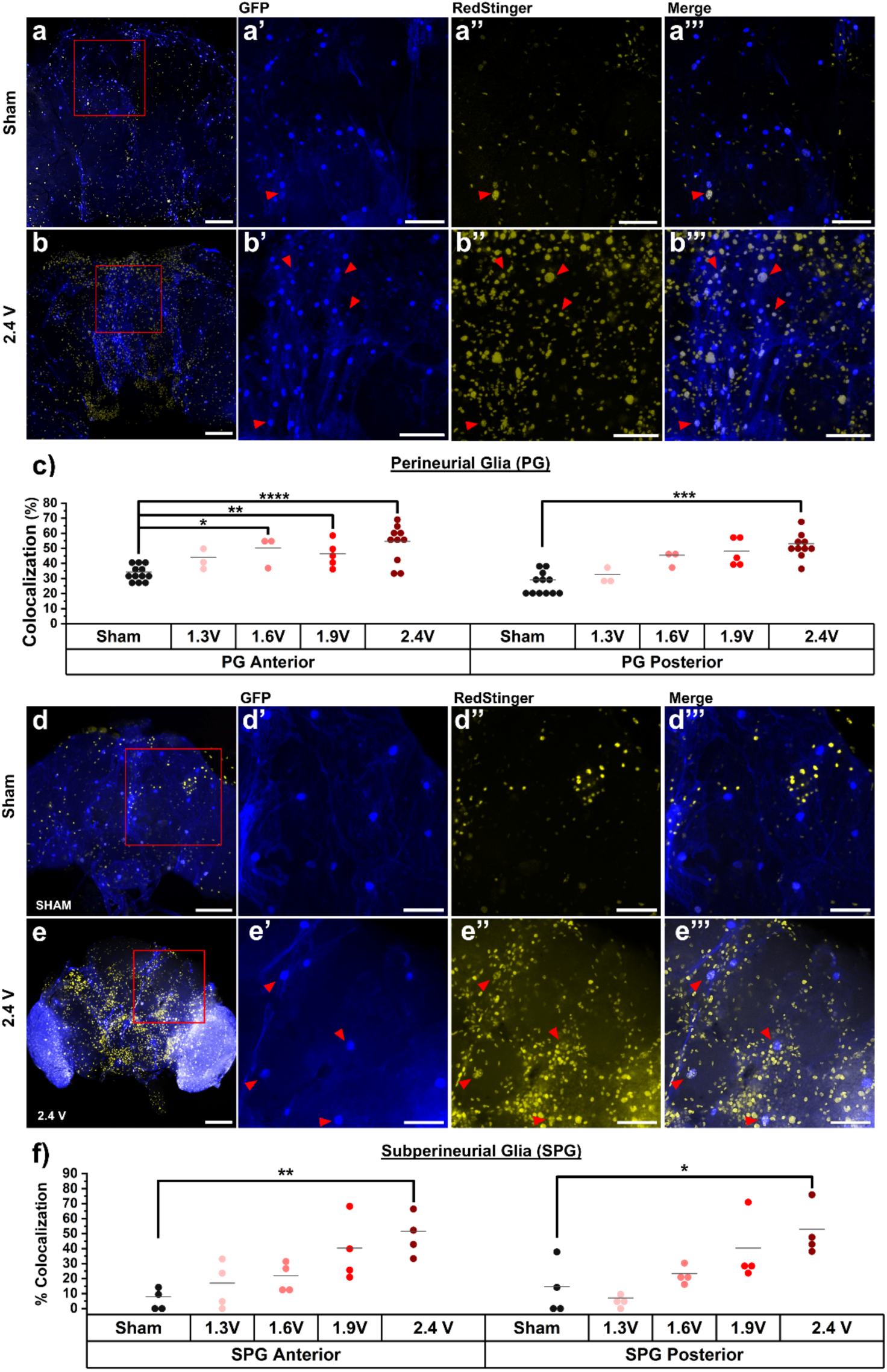
TBI severity-dependent AP-1 activation in the perineurial (PG) and subperineurial glia (SPG) of the *Drosophila* brain hemolymph-brain barrier 24 h post-injury. a-a’’’) AP-1 activation in the brains of sham control flies dissected 24 h after handling with GFP marking subperineurial glia (blue), AP-1activation marked by RedStinger expression (yellow) and colocalization shown in white. (a’-a’’’) are zoom view of red boxed region in (a). Red arrows indicate examples of colocalized nuclei. b-b’’’) As in (a-a’’’) for flies receiving 2.4 V moderate TBI. c) Quantification of colocalized RedStinger^+^/PG-GFP^+^ nuclei as a percentage of all GFP^+^ subperineurial glia nuclei at each indicated injury severity. Each symbol represents one brain. d-d’’’) As in (a-a’’’) for sham control flies with GFP marking perineurial glia. e-e’’’) As in (a-a’’’) for flies with GFP marking perineurial glia receiving 2.4 V moderate TBI. f) As in (c) for colocalized RedStinger^+^/SPG-GFP+ nuclei. (a, b, d, e) Scale bar = 100 microns (a’-a’’’, b’-b’’’, d’-d’’’, e’-e’’’) Scale bar = 50 microns. Statistical comparison by one-way ANOVA with Tukey’s post hoc test, * p<0.05, ** p<0.01, *** p<0.001, **** p<0.0001.

### Colocalization studies to identify glial subtype-specific differences in AP-1 activation following injury

While prior work in *Drosophila* has identified a pan-glial AP-1 activation response after TBI, flies have diverse glial subtypes with unique physiological roles. Using a reporter colocalization approach, we sought to determine the contribution of individual glial sub-types to the glial AP-1 activation response at 24 h post-dTBI2. Six different glia subtype Gal4 drivers (Table S1, Fig. 1) were used independently to mark distinct glial subtype nuclei with GFP (GFP.nls) in flies that have the *TRE*_*RedStinger* AP-1reporter in all cells (Pfeiffer et al. 2010; Kremer et al. 2017). Confining both fluorescent signals to the nucleus improved detection of nuclear colocalization and eliminated confounding cytoplasmic blebbing signal observed in *TRE*_*dsRed* flies post injury (Barolo et al. 2000; Saikumar et al. 2021). Dual-positive GFP.nls^+^/RedStinger^+^ nuclei represent an AP-1 activated glia of the tested glial subtype. We represent glial subtype AP-1 activation as a percentage of the number of GFP.nls^+^/RedStinger^+^ nuclei over the total number of GFP.nls^+^ glia subtype nuclei within the analyzed brain region. For all six glial subtypes tested, we compared the percentage of AP-1 activated glia between injured and sham control brains.

### Severity dependent AP-1 activation in hemolymph-brain barrier

We conducted AP-1 reporter activation colocalization studies on the two glial subclasses that comprise the *Drosophila* HBB, the PG and the SPG (Schirmeier and Klämbt 2015; Paredes-González et al. 2025). To examine AP-1 activation, we initially performed moderate (2.4 V) and sham injuries in *w^-^; TRE*_*RedStinger*/+; *PG*-*Gal4*/*UAS*-*GFP.nls* male flies and imaged fixed brains 24 h after dTBI2 injury (Fig. 2 a-b).

Consistent with other reports of PG cell morphology, many GFP^+^ nuclei were found at the periphery of the brain, in slender cells forming a loose mesh-like covering over the brain (DeSalvo et al. 2014; Kremer et al. 2017) (Fig. 2 a,a’,a’’’ and Fig. 2 b, b’,b’’’). We observed significant increases in AP-1 activation in injured flies compared to sham controls (Fig. 2 a”, b”). Computational colocalization analysis identified PG cells with activated AP-1 (a’’’, b’’’ arrowheads), quantified in Fig. 2 c. Because the HBB surrounds the periphery of the brain and each side of the brain may experience unique forces during crush injury, we analyzed the anterior and posterior sides of the HBB separately but did not find statistically significant differences in colocalization between these groups (Fig. 2 c). In sham brains, 34% ± 1.58% of GFP^+^/PG nuclei activated *TRE*_*RedStinger* expression on the anterior side and 29% ± 2.26% on the posterior side (Fig. 2c). In contrast, in flies with moderate (2.4 V) injury, 55% ± 3.80% of GFP^+^/PG nuclei activated *TRE*_*RedStinger* transcription on the anterior side and 53% ± 2.67% on the posterior side (Fig.1(Saikumar et al. 2021)1. We therefore included milder injuries (1.3 V - 1.9 V) to acquire colocalization data from more complete HBBs. These data demonstrated a trend toward injury severity-dependent increases in the number of PG-glia with activated AP-1 (Fig. 2c). These results show that 24 h after injury, activation of the AP-1 pathway occurs within PG cells on both the anterior and posterior sides of the brain, likely due to the resistant pressure of the collar to the posterior side of the head during injury.

In *w^-^; TRE*_*RedStinger*/+; S*PG*-*Gal4*/*UAS*-*GFP.nls* brains, massive polygonal shaped SPG cells were demarcated by sparse, large, oval shaped, aneuploid GFP^+^ nuclei (Fig. 2 d-d’’’, e-e’’’) (Stork et al. 2008). In sham control brains on both the anterior and posterior sides of the brain, very few SPG had active AP-1, 8% ± 4.00% and 15% ± 8.87% respectively (Fig. 2 d-d”’, 2 f). Following injuries, the percentage of colocalized nuclei trended upwards compared to shams, with only the moderate 2.4 V injury differing significantly from sham control brains on both the anterior (52% ± 7.63%) and posterior sides (53% ± 8.38%) of the brain (Fig. 2 e-e”’, Fig. 2f). These results show that, like PG cells, SPG cells significantly upregulate AP-1 transcriptional activity on both the anterior and posterior sides of the brain following dTBI2 injury, albeit with less sensitivity to low levels of injury than PG.

### AP-1 activation in cortex glia (CG) and astrocyte-like glia (ALG) following injury

In *w^-^; TRE*_*RedStinger*/+; C*G*-*Gal4*/*UAS*-*GFP.nls* brains, GFP^+^ CG nuclei were found throughout the cortex surrounding the periphery of the brain. Cytoplasmic leakage from UAS-GFP.nls revealed the honeycomb like pattern of lamellae that surround neuronal cell bodies (Fig. 3 a-a’’’, 3 b-b’’’). Whole central brain analysis of sham control brains showed very few AP-1 active cortex glia, (5% ± 1.17%) (Fig. 3 a-a’’’, c). At 24 h after 2.4 V injury, a significantly increased number of GFP^+^ CG nuclei (30% ± 6.05%) showed activation of the *TRE_RedStinger* reporter (Fig. 3 b-b’’’, c). These data show that 24 h following a moderate 2.4 V TBI, a significant proportion of cortex glia become activated.

**Figure 3.**
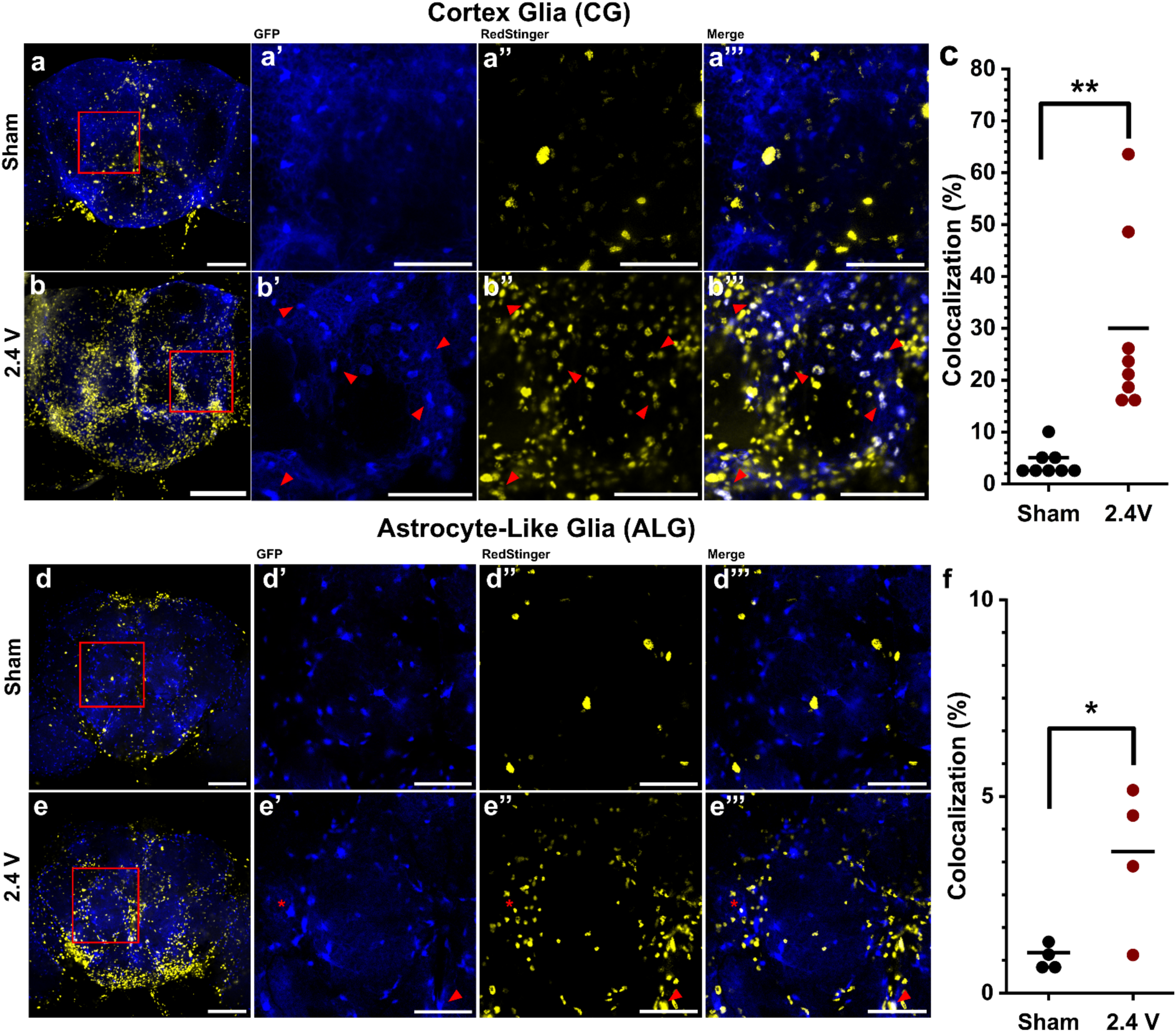
Moderate dTBI2 drives AP1 activation in cortex (CG) and little activation in astrocyte-like glia (ALG). For all image panels, brains were dissected 24 h post injury, the glial subtype is marked with GFP (blue); AP-1 activation is marked by *TRE_RedStinger* (yellow); colocalization of GFP and RedStinger (white). (a, b). *TRE_RedStinger/+*; *CG-Gal4*/*UAS-GFP.nls* (a-a’’’) Sham, (a’-a’’’) magnification of area marked by the red square in panel (a) and displayed as separate GFP, RedStinger and merged panels. Red arrows indicate examples of colocalized nuclei in all panels. Red asterisks indicate examples of overlapping, but not colocalizing, nuclei. (b-b’’’) Moderate 2.4 V injury (b’-b’’’) magnification of area marked by the red square in panel (b) and displayed as in panels a’-a’’’. (c) Quantification of colocalized RedStinger^+^/CG-GFP^+^ nuclei for sham (black) and moderate (2.4 V, dark red) injury. Each dot represents data from a colocalization analysis performed from each brain. Statistical comparison by one-way ANOVA with Tukey’s *post hoc* test. N = 8 brains per condition. (d, e) *TRE_RedStinger/+*; *ALG-Gal4*/*UAS-GFP.nls*. (d-d’’’) Sham (d’-d’’’) magnification of area marked by the red square in panel (d) and displayed as separate GFP, RedStinger and merged panels. (e-e’’’) 2.4 V injury (e’-e’’’) magnification of area marked by the red square in panel B and displayed as in panels d’-d’’. (f) Quantification of colocalized RedStinger^+^/SPG-GFP+ nuclei for sham and moderate 2.4 V injury as in (c). Each dot represents data from a whole central brain colocalization analysis. N = 4 brains per condition. (a, b, d, e) Scale bar = 100 microns (a’-a’’’, b’-b’’’, d’-d’’’, e’-e‘’’) Scale bar = 50 microns. Statistical comparison by one-way ANOVA with Tukey’s *post hoc* test. *=P>0.05, **=P>0.01.

In *w^-^; TRE*_*RedStinger*/+; *ALG*-*Gal4*/*UAS*-*GFP.nls* brains, large evenly spaced oval shaped GFP^+^ nuclei covered the neuropil surfaces (Fig. 3 d-d’’’, 3 e-e’’’). Whole brain not including optic lobe analysis of uncollared sham brains had 1% +/- 0.18% of GFP^+^ ALG nuclei which colocalized with *TRE*_*RedStinger* expression (Fig. 3 d-d’’’, 3 f). At 24 h after moderate 2.4 V injury, a small but significant increase in activated ALG cells was observed (3.6% ± 0.98%) (Fig. 3 e-e’’’, 3 f). Although the injury-induced increase in *TRE*_*RedStinger* reporter expression in ALG is statistically significant, it represents only 3.6% ± 0.98% of GFP^+^ ALG. The statistical significance likely arises from the very low baseline activation of *TRE*_*Redstinger* in this glial subpopulation.

### AP-1 activation in ensheathing glia following injury

We investigated *TRE*_*RedStinger* activation in two populations of ensheathing glia: neuropil ensheathing glia (NEG) which surround brain neuropil structures and tract ensheathing glia (TEG) that wrap axon tracts. Ensheathing glia nuclei are small (∼5 microns) and have an elongated ellipsoid shape (Kremer et al. 2017). GFP expression in NEG was driven by NEG-Gal4 (*GMR56F03*-Gal4) which drives expression in most NEG and a few TEG, while expression in TEG was driven by TEG-Gal4 (*GMR075H03*-Gal4) which drives expression in all TEG and not NEG (Kremer et al. 2017). In *w^-^; TRE*_*RedStinger*/+; *NEG*-*Gal4*/*UAS*-*GFP.nls* brains, neuropils are clearly outlined by GFP expression (Fig. 4 a-a’, b-b’). To avoid intense RedStinger signal at the brain periphery, we chose a region of interest in the interior of the brain consisting of a 15 microns thick z-stack centered around the ellipsoid body for colocalization analysis (Fig. 4 a’-a’’’, b’-b’’’). Sham control brains showed a high baseline level of *TRE_RedStinger* reporter activation in NEG (18% ± 1.24%, Fig. 4 a, a’’, a’’’, c). However, a significantly larger percentage (54% ± 5.18%) of NEG showed *TRE_RedStinger* reporter activation 24 h after dTBI (Fig. 4 b-b’’’, c). or TEG, we selected a region of interest 15 micron thick z-stack around the ellipsoid body for colocalization analysis (d-d’’’, e-e’’’). As in NEG, TEG sham control brains showed a high baseline level of TRE_*RedStinger* reporter activation (31% ± 1.82%, Fig. 4 d, d’’, d’’’, f). TEG activation of the *TRE*_*RedStinger* reporter was significantly increased 24 h after moderate dTBI, 52% ± 3.15% (Fig. 4 e, e’’, e’’’, f). To compare the extent of *TRE*_*RedStinger* reporter activation across glial subpopulations, we calculated the mean increase in reporter activation between sham and moderately injured brains 24 h after injury for each glial subtype (Fig. 5). This comparison highlights that while all glial subtypes show significant increases in reporter expression after TBI, the increase is largest and most consistent in the NEG (36% ± 5.33%), Fig. 5 b). While a similar average increase is seen in CG, CG also showed more variable reporter responses at baseline resulting in a larger standard error compared to controls (25% ± 6.16%). PG and TEG also show highly consistent increases in the percentage of cells activating the *TRE*_*Redstinger* reporter, albeit smaller than the NEG.

**Figure 4.**
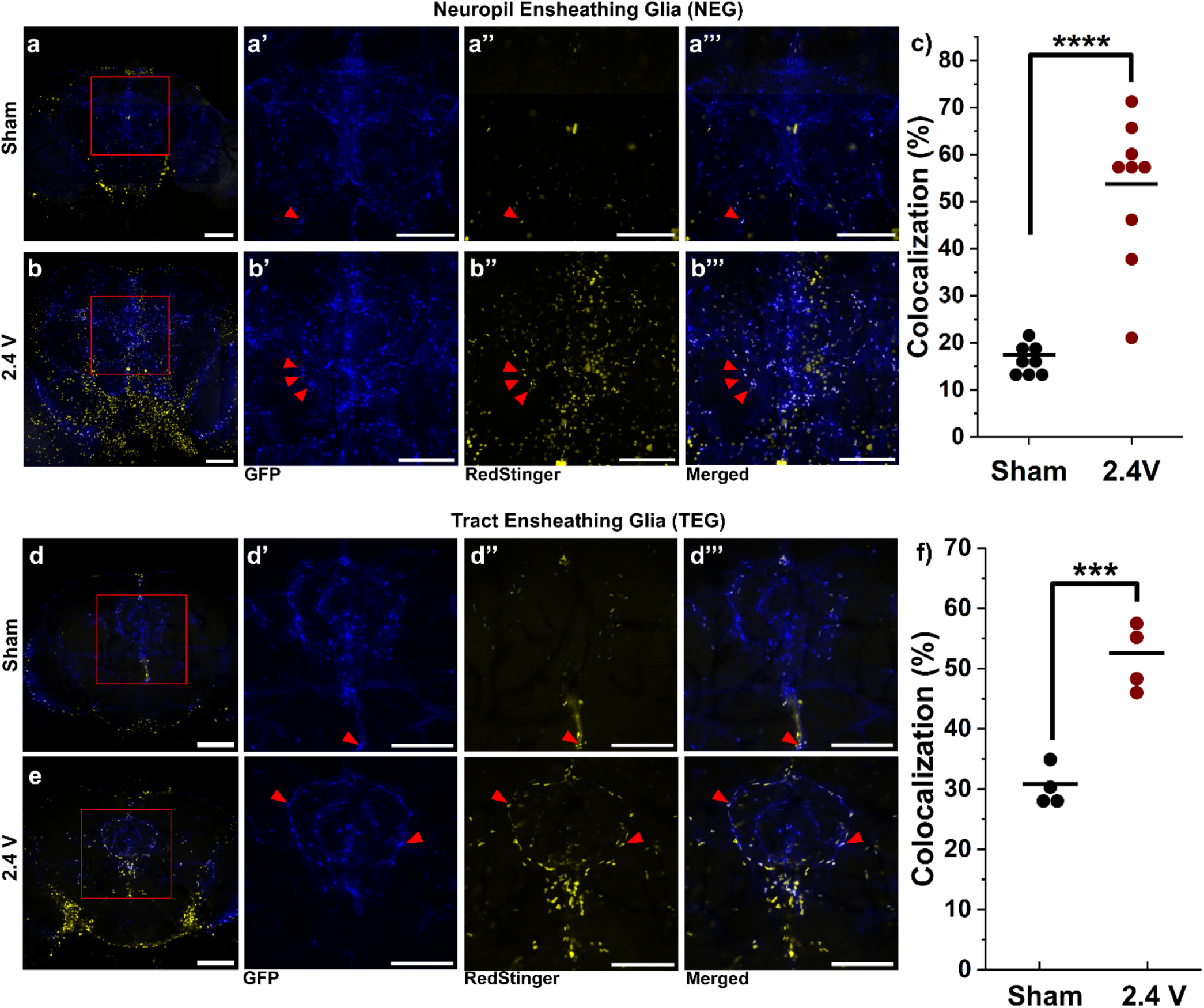
Moderate dTBI2-drives AP1 activation in neuropil (NEG) and tract ensheathing glia (TEG). For all image panels, brains collected 24 hours post injury, glial subtype is marked with GFP (blue); AP-1 activation marked by *TRE_RedStinger* (yellow); colocalization of GFP and RedStinger (white). (a, b) *TRE_RedStinger/+*; *NEG-Gal4*/ *UAS-GFP.nls*. (a-a’’’) Sham (a’-a’’’) magnification of area marked by the red square in panel (a) and displayed as separate GFP, RedStinger and merged panels. Red arrows indicate colocalized nuclei. (b-b’’’) moderate 2.4 V injury (b’-b’’’) magnification of area marked by the red square in panel (b) and displayed as in panels (a’-a’’’). (c) Quantification of colocalized RedStinger^+^/NEG-GFP^+^ nuclei between moderate 2.4 V injury (red) and sham (black). Each dot represents data from a colocalization analysis performed from a region centered around the ellipsoid body. N= 4 brains per condition. (d, e) *TRE_RedStinger*; *TEG-Gal4*/ *UAS-GFP.nls*. (d-d’’’) Sham (d’-d’’’) magnification of area marked by the red square in panel D and displayed as in panels (a’-a’’’). (e-e’’’) 2.4 V injury (e’-e’’’) magnification of area marked by the red square in panel (e) and displayed as in panels (a’-a’’’). (f) Quantification of colocalized *TRE_RedStinger+/TEG-GFP+* nuclei at sham and moderate 2.4 V injury as in (c). Each dot represents data from a colocalization analysis performed from a region centered around the ellipsoid body. N = 8 brains per condition. (a, b, d, e) Scale bar = 100 microns (a’-a’’’, b’-b’’’, d’-d’’’, e’-e‘’’) Scale bar = 50 microns. Statistical comparison by one-way ANOVA with Tukey *post hoc* test, ***=P>0.001, ****=P>0.0001.

**Figure 5.**
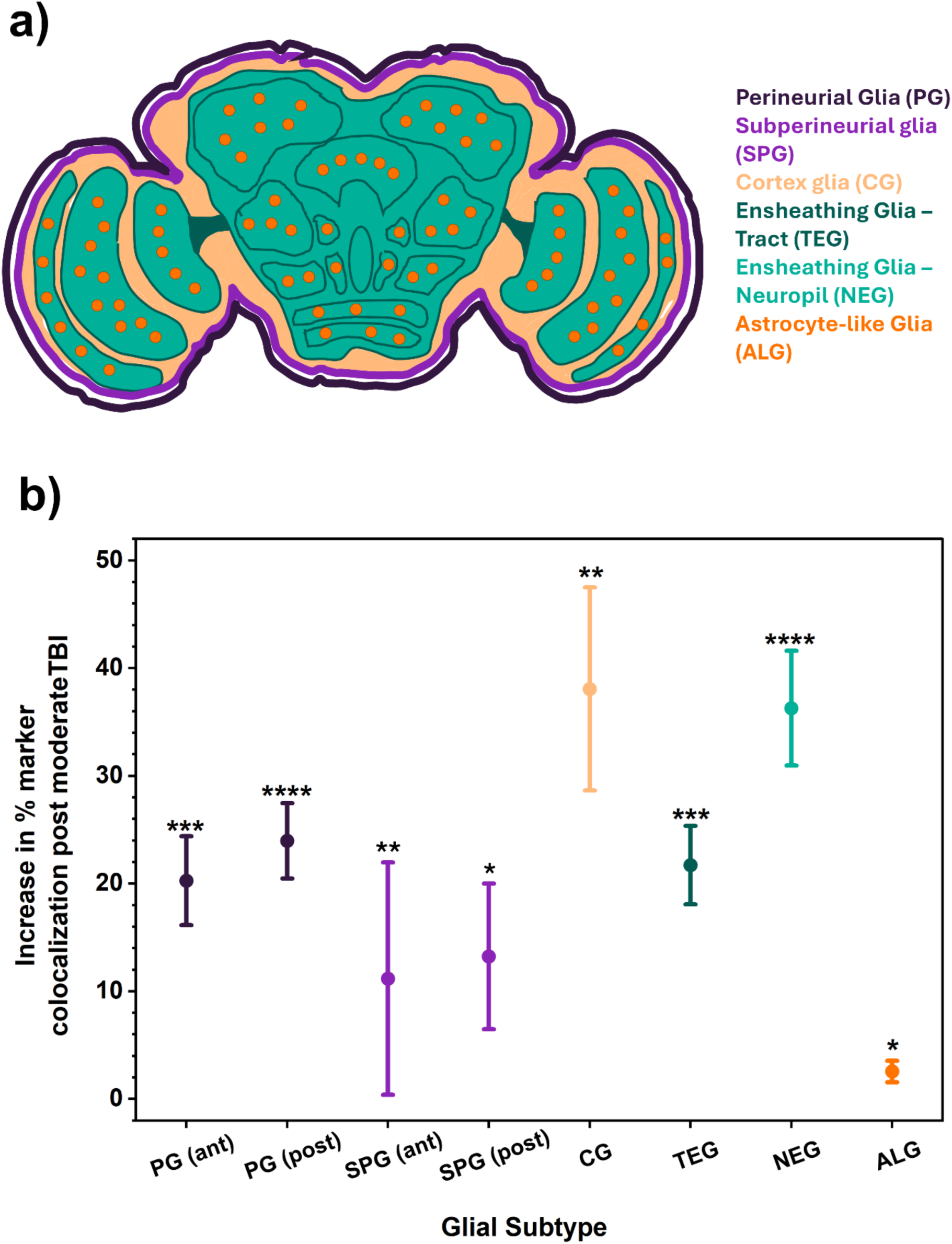
TBI induces heterogeneous AP-1 activation levels across glial subtypes. a) Schematic depiction of glial subtypes in the *Drosophila* brain. Perineurial glia (dark purple) and subperineurial glia (purple) make up the outer and inner layers of the blood-hemolymph-brain barrier respectively. Cortex glia (peach) surround neuronal cell bodies. Tract ensheathing glia (dark green) wrap axonal tracts between neuropils while neuropil ensheathing glia (teal) wrap neuropil borders. Astrocyte-like glia (orange) are located throughout the neuropil. b) Summary comparison of the increase in percentage of cells of each glial subtype showing colocalization with the AP-1 *TRE*_*RedStinger* reporter 24 h after moderate injury. Statistical significance was determined by one-way ANOVA of the percent colocalization for each individual brain or brain region for each glial subtype as shown in plots in Figs. 2-4. * p<0.05, ** p<0.01, *** p<0.001, **** p<0.0001.

Many AP-1 positive nuclei observed after injury do not overlap with pan-glial *repo*-GAL4^M1B^ or with pan-neuronal *nSyb*-Gal4 marker expression (Fig. S2 a, b). We hypothesized that the unidentified nuclei may be macrophages recruited to areas of injury (Zechini et al. 2025). To test this, we used the *He*-Gal4 driver to label macrophages and test colocalization with AP-1 activation after dTBI2. However, in *w^-^; TRE*_*RedStinger*/+; *He*-*Gal4*/*UAS*-*GFP.nls* brains 24 h after moderate (2.4 V) injury, only a few GFP^+^ macrophage nuclei were observed at the periphery of the brain but not in the central brain (Fig. S2 c-c’’’). As *He*-Gal4 is expressed in the majority of macrophages, it is unlikely that unidentified AP-1 positive nuclei are macrophages, and thus the identity of these cells remains to be determined (Makhijani et al. 2011).

### Pan-glial inhibition of AP-1 by Kay^DN^ reduces post injury survival

Prior studies have found that inducible pan-glial genetic inhibition of AP-1 signaling reduces fly survival after TBI (Byrns et al. 2021). We used a similar approach to examine the effects of up- and downregulation of glial AP-1 signaling on survival following dTBI2. For initial experiments, we used the pan-glial driver *repo*-GAL4^M1B^ to overexpress either an activating or inhibiting transgene of the AP-1 pathway (Fig. 6 a, b). To activate the AP-1 pathway, we overexpressed a constitutively active mutant of *rolled* (*rl*), the fly homolog of ERK MAP kinase (UAS-*rl*^R80S+D334N^) (Kushnir et al. 2020). *Drosophila* Rolled phosphorylates and activates the D-Fos, D-Jun AP-1 transcription factor heterodimer, and is activated following *Drosophila* head injury (Byrns et al. 2021). To inhibit AP-1 signaling, we expressed the dominant negative form of *kayak* (D-Fos). *kay*^DN^ expresses only the bZIP domain of D-Fos without the DNA binding motif and is able to bind to other bZIP containing transcription factors and render them transcriptionally inactive.

**Figure 6.**
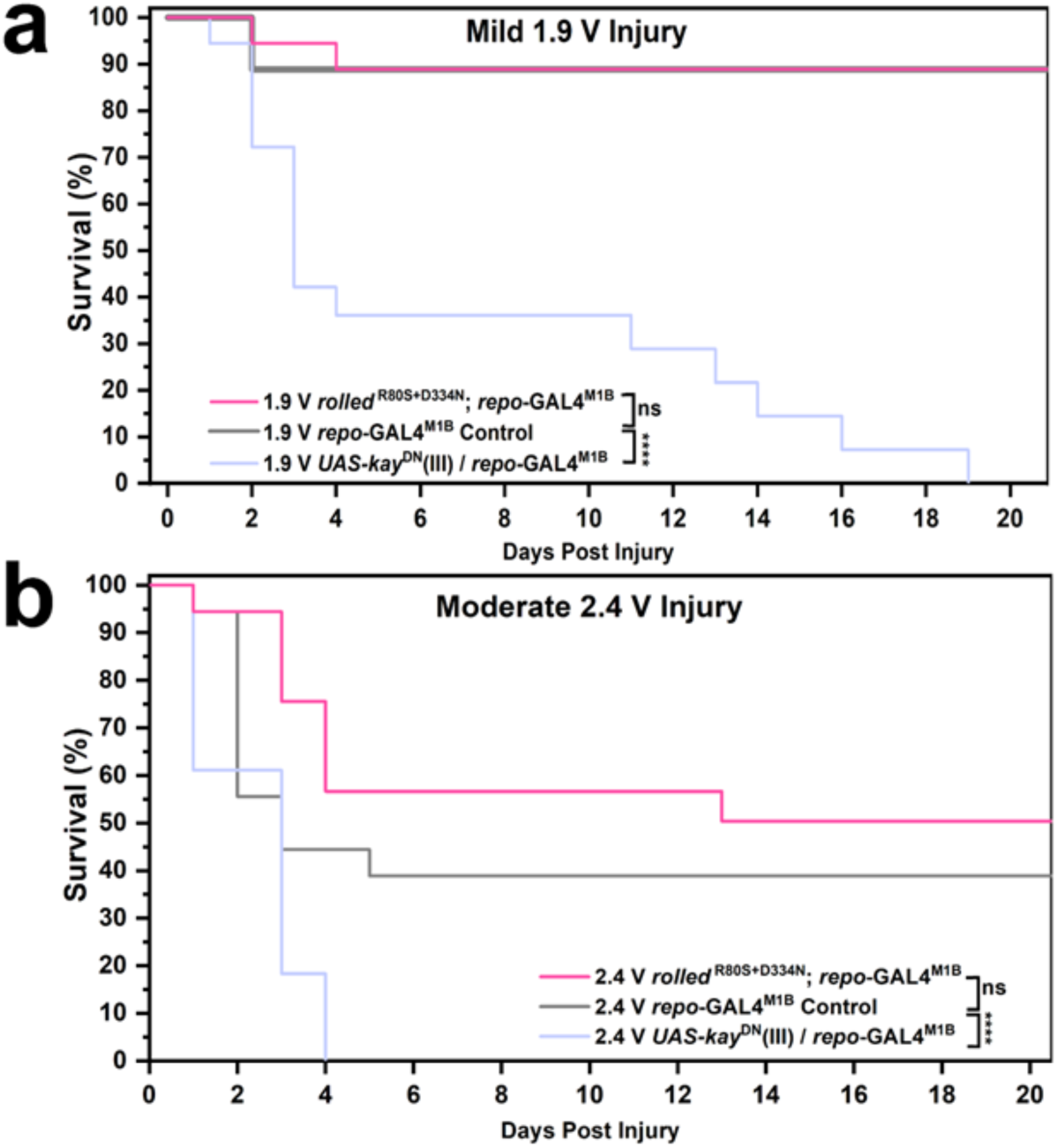
Pan-glial genetic inhibition of AP-1 signaling reduces post-dTBI2 survival. Twenty-one day survival was analyzed following a) Mild 1.9 V injury and b) Moderate 2.4 V injury to flies carrying the pan-glial *repo*-*Gal4*^M1B^ driver with either the AP-1 activator (*rolled*^R80S+D334N^/+; *repo*-*Gal4*^M1B^/+, red), the AP-1 repressor (*UAS*-*kay*^DN^ (III) /+ ; *repo*-*Gal4*^M1B^/+, blue) and a Gal4 control (*repo*-*Gal4*^M1B^/+, dark gray). Log-rank pairwise comparison was used to determine statistical difference between injured groups. N = 18 flies per condition, one biological replicate. Brackets in legend designate the statistical significance between survival curves, ns =not significant, **** p<0.0001.

We performed mild (1.9 V) and moderate (2.4 V) injuries on three-day old *w^-^;* UAS-*rl*^R80S+D334N^/+; *repo*-GAL4^M1B^/+ and *w^-^;* UAS-*kay*^DN^ (III)/*repo*-GAL4^M1B^ male flies and compared survival to the Gal4 only parental control (*w^-^; repo*-GAL4^M1B^/+) (Fig. 6 a, b). Injured flies expressing *rl*^R80S+D334N^ in all glia did not show significant differences in survival compared to the Gal4 parental control after mild or moderate injury. However, pan-glial inhibition of the AP-1 pathway using *kay*^DN^ caused a significant increase in mortality following injuries. Following a mild 1.9 V injury, more than half of *w^-^;* UAS-*kay*^DN^ (III)/*repo*-GAL4^M1B^ flies died by three days post-injury (dpi) and all perished by 19 dpi. In controls, ∼90% of injured Gal4 parental control flies survived past 21 dpi.

Following the 2.4 V injury, all *w^-^;* UAS-*kay*^DN^ (III)/*repo*-GAL4^M1B^ flies died by 4 dpi whereas 40% of injured Gal4 parental control flies survived past 21 dpi. These results show that expression of *kay*^DN^ in all glia severely increases lethality following injury.

Pan-glial expression of two different UAS-*kay*^DN^ fly lines showed different responses to sham handling (Fig. S3 a-d). Using the second chromosome line UAS-*kay*^DN^ (II), we found that *w^-^;* UAS-*kay*^DN^ (II)/+; *repo*-GAL4^M1B^/+ uninjured sham flies die by 3 dpi (Fig. S3 a, c). Flies survive longer using the third chromosome UAS-*kay*^DN^ (III) line as 50% of uninjured sham *w^-^;* UAS-*kay*^DN^ (III)/ *repo*-GAL4^M1B^ survive past 21 dpi (Fig. S3 b, d). Because of the fragility of the *w^-^;* UAS-*kay*^DN^ (II)/+; *repo*-GAL4^M1B^/+ flies to even sham handling, these flies showed significantly reduced mortality compared to parental control lines, as there was no significant mortality difference between sham and mild injury within this genotype (Fig. S3a). However, after moderate injury, the *w^-^;* UAS-*kay*^DN^ (II)/+; *repo*-GAL4^M1B^/+ flies showed significantly reduced mortality to both parental controls and their own sham controls. As discussed above, UAS-*kay*^DN^ (III)/ *repo*-GAL4^M1B^ flies show a significant increase in mortality following mild or moderate head injury compared to both Gal4 and UAS controls and to its own sham controls (Fig. 6; Fig. S3 b, d). These results suggest that the second chromosome (II) UAS-*kay*^DN^ is expressed more strongly than the third chromosome transgene. All further survival experiments utilized the UAS-*kay*^DN^ (II) construct.

### Ensheathing glia-specific activation of AP-1 is necessary for acute TBI survival

We found that inhibiting the AP-1 pathway in neuropil ensheathing glia reduces survival following dTBI2 (Fig. 7 a-b’). Using the same glia subtype *Gal4* drivers used for brain imaging analysis, we inhibited AP-1 by expressing UAS-*kay*^DN^ (II) in the PG, SPG, CG, TEG, NEG and ALG glia subtypes. We injured these flies with a mild 1.9 V injury and compared survival to the *w^-^;* UAS-*kay*^DN^ (II)/+ control. We found that only *w^-^;* UAS-*kay*^DN^ (II)/+; NEG-Gal4/+ flies have significantly reduced survival following injury compared to the UAS control (Fig. 7 a).

**Figure 7.**
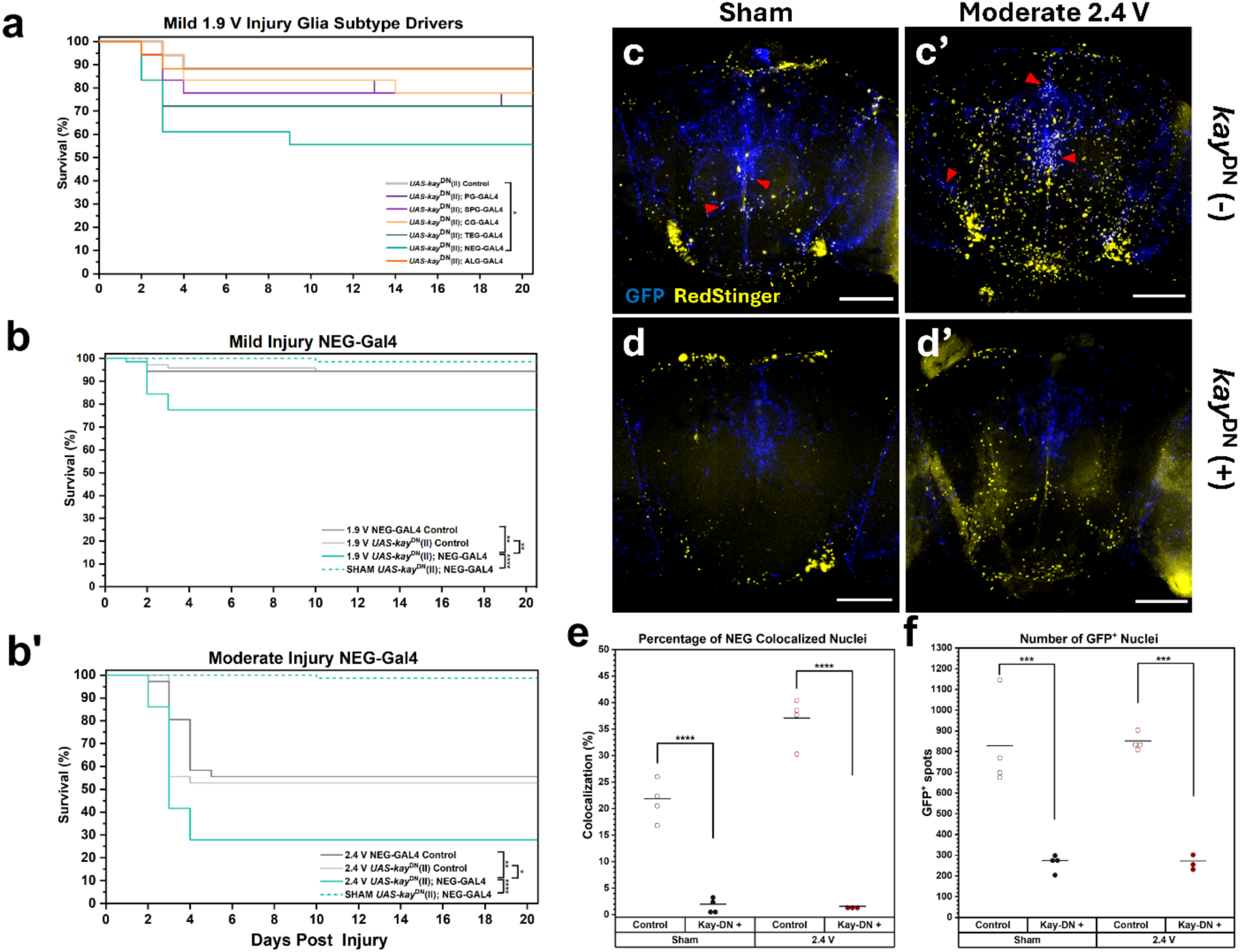
Neuropil ensheathing glia-specific inhibition of AP-1 signaling reduces post-dTBI2 survival and NEG cell numbers. a) Twenty-one day survival was analyzed following a Mild 1.9 V injury to *UAS*-*kay*^DN^ (II) /+ control (light gray) and *UAS*-*kay*^DN^ (II) /+; *Glia subtype*-Gal4/+ flies; PG (purple), SPG (light purple), CG (yellow), TEG (dark green), NEG (light green), ALG (orange). N = 18 flies per condition. All injuries were performed on the same day. b, b’) Large scale twenty-one day survival was analyzed following a (b) mild 1.9 V injury and a (b’) moderate 2.4 V injury to *NEG-Gal4*/+ control (dark gray), *UAS*-*kay*^DN^ (II) /+ control (light gray), *UAS*-*kay*^DN^ (II) /+; *NEG-Gal4*/+ (solid light green) or Sham *UAS*-*kay*^DN^ (II) /+; *NEG-Gal4*/+ (dashed light green). (b) N=72 flies per condition; (b’) N=36 flies per condition. All injuries were performed on the same day. Log-rank pairwise comparison was used to determine statistical difference between injured groups for survival assays. c, c’) *TRE_RedStinger/+*; *NEG-Gal4*/*UAS-GFP.nls* brains imaged for GFP (blue) and RedStinger (yellow) expression. c) Uncollared Sham (c’) moderate 2.4 V injury. d, d’) *TRE_RedStinger/ UAS*-*kay*^DN^ (II); *NEG-Gal4*/*UAS-GFP.nls* brains imaged for GFP and RedStinger expression. (d) Uncollared Sham (d’) moderate 2.4 V injury. All brains dissected 24 hours post injury. e) Quantification of colocalized RedStinger^+^/NEG-GFP^+^ nuclei between Sham brains without *UAS*-*kay*^DN^ (II) (empty black circle) or with *UAS*-*kay*^DN^ (II) (filled black circle) and moderate 2.4 V injury brains without *UAS*-*kay*^DN^ (II) (empty red circle) or with *UAS*-*kay*^DN^ (II) (filled red circle). N= 4 brains per condition, except moderate 2.4 V injury brains with *UAS*-*kay*^DN^ (II) N=3. (e) Quantification of colocalized RedStinger^+^/NEG-GFP^+^ nuclei between Sham brains without Kay^DN^ (-) (empty black circle) or with Kay^DN^ (+) (filled black circle) and moderate 2.4 V injury brains without Kay^DN^ (-) (empty red circle) or with Kay^DN^ (+) (filled red circle). N= 4 brains per condition, except moderate 2.4 V injury brains with Kay^DN^ (+) N=3. (f) Quantification of number of NEG/GFP+ nuclei using the same genotypes, conditions, brains and labels as used in (e). e, f) Each dot represents data from analysis performed on central brain sections excluding the optic lobes starting from the most anterior to just posterior to the ellipsoid body. Statistical comparison by one-way ANOVA with Tukey’s *post hoc* test. Brackets in legend and graphs designate the statistical significance between data sets, * p<0.05, ** p<0.01, *** p<0.001, **** p<0.0001. Scale bar = 100 microns (c, c’, d, d’).

To follow-up, we performed a large-scale experiment comparing the survival rate of injured *w^-^;* UAS-*kay*^DN^ (II)/+; NEG-Gal4/+ flies to the injured *w^-^;* UAS-*kay*^DN^ (II)/+ control, injured *w^-^;* NEG-Gal4/+ control and the sham *w^-^;* UAS-*kay*^DN^ (II)/+; NEG-Gal4/+ (Fig. 7 b, b’). Following either a mild 1.9 V injury or a moderate 2.4 V injury, *w^-^;* UAS-*kay*^DN^ (II)/+; NEG-Gal4/+ flies show a significant increase in mortality compared to both the injured UAS and Gal4 controls. Inhibition of AP-1 in NEG throughout development and in the adult fly does not compromise viability, as nearly 100% of sham *w^-^;* UAS-*kay*^DN^ (II)/+; NEG-Gal4/+ flies survive past 21 dpi.

When *kay*^DN^ is expressed in NEG, we saw a significant decrease in the number of NEG cells per brain in both the shams, 445 ± 61 and injured brains, 446 ± 37 (Fig. 7 d, d’, f).

## DISCUSSION

Deciphering the cellular and molecular genetic responses to brain injury provides critical insight towards defining the steps required for full brain recovery. In both mammals and insects, it is well documented that after injury, glia participate in nervous system recovery (Sofroniew and Vinters 2010). After injury, *Drosophila* glia activate a massive AP-1 transcriptional response that is required for survival (Byrns et al. 2021). We used a second-generation head-specific TBI paradigm and found that glial AP-1 responses scaled with injury severity (Fig. 1). We further defined the *Drosophila* AP-1 injury induced glia response by profiling AP-1 activation after dTBI across glial subpopulations (Figs. 2-4). This approach identified NEG as having particularly large increase in AP-1 activation 24 hpi (Fig. 5). We tested genetic approaches to activate or inhibit AP-1 and found that inhibition of AP-1 signaling by overexpression of Kay^DN^ resulted in reduced survival after mild or moderate injury (Fig. 6). Genetic inhibition of AP-1 signaling in each glial sub-population identified NEG as a key cell type where AP-1 signaling is required for survival after mild or moderate dTBI (Fig. 7). We also found that genetic inhibition of AP-1 in NEG reduced the number of NEG cells in the adult brain, which may indicate that the presence of sufficient numbers of NEG are necessary for survival after injury.

Using colocalization studies, we observed strong and consistent AP-1 activation in multiple glia subtypes after injury. Our study revealed the importance of nuclear fluorescence labeling to clearly visualize the TBI-induced AP-1 response *in vivo*. Our colocalization image analysis was able to determine if AP-1 was “on” or “off” within individual cells. Improved image analysis can attempt to better determine fluorescent intensity levels of the AP-1 response, as transcriptional intensity greatly affects cellular outputs (Volteras et al. 2024). Surprisingly, many TBI induced AP-1 activated nuclei did not overlap with glia markers and lacked a corresponding Gal4 driver line for further genetic manipulation studies. Nuclear labeled cell sorting of injured brains coupled with RNAseq can help to further define the AP-1 cellular response and uncover additional genetic tools which can be used to study the TBI response at the cellular and molecular levels. Interestingly, in the PG, TEG, and NEG glia which activate AP-1 after injury, we observed low levels of AP-1 activation in sham brains. This shows that even in uninjured brains, a proportion of cells possess the cellular machinery to activate AP-1 and perhaps AP-1 activation serves to amplify an already occurring cellular function or property in which PG, TEG and NEG are involved prior to injury.

To test the role of post-TBI glial AP-1 activation in survival, we activated and inhibited the AP-1 pathway by overexpressing Rolled^R80S+D334N^ and Kay^DN^, respectively, in all glia cells and in each glial subtype. Pan-glial overexpression of constitutively active Rolled^R80S+D334N^ had no effect on survival following injury (Fig. 6). However, we find that constitutive pan-glial inhibition of the AP-1 pathway by overexpression of UAS-*kay*^DN^ causes lethality in uninjured adult flies. This is consistent with prior observation that temporally inducible pan-glial inhibition of AP-1 signaling using the GeneSwitch system also significantly reduces survival of sham flies compared to vehicle controls (Byrns 2021). However, when AP-1 is inhibited within specific glial subtypes by UAS-*kay*^DN^ overexpression, uninjured flies survive past the 21-day test period. These results suggest that functional AP-1 signaling is required in multiple glial subtypes for normal development and lifespan in uninjured flies.

After head injury, inhibition of AP-1 in all glia and specifically in the NEG subtype, causes an increase in mortality compared to genetic controls. *Drosophila* ensheathing glia are known to migrate to axonal injury and actively phagocytose cellular debris, which requires activation of AP-1 via the CED-1 receptor, Draper (Lu et al. 2017; Purice et al. 2017). However, others have shown that increased phagocytosis following head injury occurred normally in *draper*^−/−^ mutants and *draper-RNAi* had no effect on AP-1 activation following head injury as monitored by the *TRE*_*dsRed* reporter (Byrns et al. 2021). Further experiments are needed to determine if phagocytosis by NEG cells is required for optimal survival and if upregulation of phagocytosis requires AP-1 activation following injury. Interestingly, AP-1 inhibition in NEG but not TEG increased mortality following injury. Possible explanations for this observation could arise due to putative differences in Gal4 expression levels between Gal4 drivers or previously described differences in the cell number and location of NEG and TEG-Gal4 expressing cells (Kremer et al. 2017).

We found that NEG-specific overexpression of Kay^DN^ in NEG cells resulted in fewer NEG cells present in the adult brain compared to WT controls. Others have shown that the number of NEG cells declines with age in both ants and *Drosophila,* and that expression of the anti-apoptotic gene *p*35 in NEG rescued the age-related decline in cell number, improved the neuromotor performance of aged flies, and extended lifespan (Sheng et al. 2023). AP-1 is implicated in both pro- and anti-apoptotic processes, depending on cell type and environmental context (Ameyar et al. 2003; Hess et al. 2004). AP-1 activity has been associated with pro-apoptotic transcription through Jun kinase (JNK) mediated signaling during development (Adachi-Yamada et al. 1999;

Takatsu et al. 2000; Lehmann et al. 2002). However, AP-1 is known to interact with a nuclear receptor transcriptional silencing complex subunit, Ebi, and suppress basal transcription levels of pro-apoptotic genes in sensory neurons (Lim et al. 2012). AP-1 activity is also linked with aging-associated glia senescence, suggesting that TBI-driven induction of glial AP-1 activity may accelerate aging-related pathologies (Martínez-Zamudio et al. 2020; Byrns et al. 2024). The potential role of senescent glia after injury is unclear, as senescent cells promote wound healing but are also pro-inflammatory which could worsen tissue decline. Which glial subtypes enter senescence during normal aging, and whether other markers of glial senescence are induced by TBI remain to be determined.

We have shown that AP-1 activation within NEG cells increases after injury. In addition, constitutive inhibition of AP-1 signaling within NEG reduces the number of NEG in the brain which may lead to the observed increase in mortality following TBI. Our findings validate the utility of the *Drosophila* dTBI paradigm for investigating the contributions of glial subtypes to post-injury physiology and pathology. Our data provides compelling evidence for further investigation of the downstream molecular consequences of AP-1 activation in NEG, including roles in cellular development and survival. In addition, the phagocytic nature of NEG suggests the possibility that AP-1 activation within NEG affects its ability to clear debris in response to TBI which may affect survival. Apart from NEG, our studies show that multiple glial subtypes become activated after TBI. It is possible that multiple glial subtypes act synergistically for brain recovery through AP-1 signaling, as demonstrated by the severe TBI mortality outcomes found when inhibiting AP-1 activation pan-glially. This can be further investigated by inhibiting AP-1 signaling pairwise in glia subtypes to identify populations with synergistic or antagonistic effects on fly survival or other post-TBI outcomes. As *Drosophila* possess many similar cell types and molecular signaling pathways found in humans, determining the underlying cellular and molecular genetics of fly TBI recovery can provide useful comparative information toward improving assessment and treatment of human TBI.

## DATA AVAILABILITY STATEMENT

Fly strains and plasmids are available upon request. All data necessary to replicate plots and statistical analysis are available on Zenodo, DOI: 10.5281/zenodo.21839503

## ACKNOWLEDGEMENTS

Image acquisition and analysis was made possible by the Waksman Institute Shared Imaging Core Facility, and The Human Genetics Institute Imaging Core Facility at Rutgers. Thanks to Dr. Nanci Kane for her expert assistance with image acquisition and to Dr. Bipin Tripathi for assistance with the smurf assay. Thanks to members of the Barber, Irvine, and McKim labs for helpful discussion.

Thanks to Ze’ev Paroush, Lynn Cooley, and Achim Paululat for reagents. Thanks to the Bloomington *Drosophila* Stock Center and Flybase for their service to the fly community. Stocks obtained from the Bloomington *Drosophila* Stock Center (NIH P40OD018537) were used in this study.

## SUPPLEMENTARY TABLE AND FIGURE LEGENDS

**Table S1. Reagent Table.** Chemicals, fly stocks, plasmids and software used in this study.

**Table S2. PCR Primers and templates used to generate *pH_TRE_RedStinger_attB***. PCR products, primers and templates used to generate *pH_TRE_RedStinger_attB*. Primers are written in the 5’ to 3’ direction. Generic Barber lab names are provided for PCR fragments and for primer names found in parenthesis. Genomic DNA was from *TRE*_*dsRed* transgenic flies.

**Table S3. Substack parameters by glial subtype.** Whole brain images were made into substacks prior to Imaris colocalization analysis which differed depending upon the glia subtype being examined. The number, location and the thickness of the substacks used for different glia subtypes is listed.

**Table S4. Spot XYZ parameters for GFP by glial subtype**. To reduce erroneous spot assignment to background or tissue abnormalities, Imaris spot assignment to GFP^+^ nuclei was limited to parameters that correspond to the expected nuclei size range of the tested glial subtype.

**Table S5. Number of GFP^+^ glia subtype spots used for colocalization analysis per area analyzed.** Number of GFP^+^ glia subtype spots tested for colocalization with RedStinger differed depending upon which glia subtype was being tested.

**Figure S1. dTBI2-induced mortality is not due to gut barrier leakage.** Flies were fed a less nutritive dyed diet beginning 2 days prior and ending 24 h post-injury. a) Representative male fly subjected to moderate 2.4 V injury using dTBI2, showing no smurfing as dye is consolidated in the gut. b) Representative male fly subjected to moderate TBI using mixer mill at 30 seconds and 100 rpm, showing smurfing as gut barrier integrity is disrupted leading to dye leaking throughout the body. c) Table of experimental information and percent of flies with gut barrier disruption post-injury using mixer mill only.

**Figure S2. Non-glial and non-neuronal cells show AP-1 activation after TBI.** For all image panels, brains were dissected 24 h post injury, GFP (blue) and RedStinger (yellow), colocalization of GFP and RedStinger (white). a) Moderate 2.4 V injury, *TRE_RedStinger/+*; *repo*-*Gal4*^M1B^/*UAS-GFP.nls.* b) Mild 1.9V injury, *TRE_RedStinger/+*; *nSyb*-Gal4/*UAS-GFP.nls*, (a, b) Single serial confocal sections of a single brain arranged from the antennal anterior side (left) to the posterior side (right) of the brain. (c, c’, c’’, c’’’) *TRE_RedStinger/+*; *He-Gal4*/*UAS-GFP.nls*, moderate 2.4 V injury, magnification of area marked by the red square in panel (c) and displayed as separate (c’) GFP, (c”) RedStinger and (c’’’) merged panels. Red arrows indicate examples of colocalized nuclei in all panels. (a,b,c) Scale bar = 100 microns (c’,c’’,c’’’) Scale bar = 50 microns.

**Figure S3. Pan-glial expression of UAS-*kay*^DN^ shows variable mortality phenotypes depending upon chromosomal location of transgene.** Twenty-one day survival experiments examining effects of injury on flies expressing either *UAS*-*kay*^DN^ (II) or *UAS*-*kay*^DN^ (III) by the pan-glia *repo*-*Gal4*^M1B^ driver. (a, b) mild 1.9 V injury (c, d) moderate 2.4 V injury. (a, b, c, d) *repo*-*Gal4*^M1B^/+ control (dark gray), (a, c) *UAS*-*kay*^DN^ (II) /+ control (light gray), (b, d) *UAS*-*kay*^DN^ (III) /+ control (light gray), (a, c) *UAS*-*kay*^DN^(II) /+; *repo*-*Gal4*^M1B^ /+ experimental (black), (b, d) *repo*-*Gal4*^M1B^ /*UAS*-*kay*^DN^ (III) experimental (black), (a, b, c, d) Experimental Shams (dashed black). N = 18 flies per condition. For all survival experiments, Sham flies were collared, left in the collar several minutes then removed with forceps. Log-rank pairwise comparison was used to determine statistical differences between injured groups for survival assays. Brackets in legend and graphs designate the statistical significance between data sets, ns = not significant, * p<0.05, ** p<0.01, *** p<0.001, **** p<0.0001.

