## Supplementary Information for "AP-1 activation in *Drosophila* neuropil ensheathing glia improves traumatic brain injury survival"

### SUPPLEMENTARY MATERIAL

| **Table S1: Reagent Table** | | | |
|---|---|---|---|
| **Reagent type or resource** | **Designation** | **Source or Reference** | **Identifiers** |
| Chemicals, peptides, and recombinant proteins | | | |
| Chemical compound | Erioglaucine disodium salt | Millipore Sigma 861146 | CAS Number: 3844-45-9 |
| Enzymes | NEBuilder® | New England Biolabs | E2621L |
| Chemical compound | Vectashield® | VectorLabs | H-1000-10 |
| Experimental models: Organisms/strains | | | |
| Genetic reagent (D. melanogaster) | *Iso*$^{31}$ | Bloomington *Drosophila* Stock Center (Ryder et al. 2004) | BDSC: 5905 |
| Genetic reagent (D. melanogaster) | *TRE_dsRed* | Bloomington *Drosophila* Stock Center | BDSC: 59011 FLYB: FBst0059011 |
| Genetic reagent (D. melanogaster) | *TRE_RedStinger* | GMR54C07-GAL4 | |
| Genetic reagent (D. melanogaster) | *repo*-GAL4$^{M1B}$ | (Jefferis et al. 2004) | FLYB: FBal0154701 |
| Genetic reagent (D. melanogaster) | *GMR85G01*-GAL4 PG-Gal4 | Bloomington *Drosophila* Stock Center | BDSC:40436; FLYB: FBti0139096 |
| Genetic reagent (D. melanogaster) | *GMR54C07*-GAL4 SPG-Gal4 | Bloomington *Drosophila* Stock Center | BDSC: 50472; FLYB: FBti0136765 |
| Genetic reagent (D. melanogaster) | *GMR54H02*-GAL4 CG-Gal4 | Bloomington *Drosophila* Stock Center | BDSC: 45784; FLYB: FBti0136820 |
| Genetic reagent (D. melanogaster) | *GMR86E01*-GAL4 ALG-Gal4 | Bloomington *Drosophila* Stock Center | BDSC: 45914 FLYB: FBti0139155 |
| Genetic reagent (D. melanogaster) | *GMR56F03*-GAL4 NEG-Gal4 | Bloomington *Drosophila* Stock Center | BDSC: 39157 FLYB: FBti0136979 |
| Gene (*D. melanogaster*) | *GMR075H03*-GAL4 TEG-Gal4 | Bloomington *Drosophila* Stock Center | BDSC: 39908 FLYB: FBti0138281 |
| Genetic reagent (*D. melanogaster*) | UAS-*GFP.nls* (II) | Bloomington *Drosophila* Stock Center | BDSC: 4775 FLYB: FBti0012493 |
| Genetic reagent (*D. melanogaster*) | UAS-*GFP.nls* (III) | Bloomington *Drosophila* Stock Center | BDSC: 4776 FLYB: FBti0012493 |
| Genetic reagent (*D. melanogaster*) | UAS-*rl*$^{R80S+D334N}$ | (Kushnir et al. 2020) Gift of Ze'ev Paroush | FLYB: FBal0358599 |
| Genetic reagent (*D. melanogaster*) | UAS-*kay*$^{DN}$ (II) | Bloomington *Drosophila* Stock Center | BDSC: 7214 FLYB: FBti0228767 |

| | | | |
|---|---|---|---|
| Genetic reagent (*D. melanogaster*) | UAS-*kay*$^{DN}$ (III) | Bloomington *Drosophila* Stock Center | BDSC: 7215<br>FLYB: FBti0228767 |
| Genetic reagent (D. melanogaster) | *nSyb*-Gal4 | Bloomington *Drosophila* Stock Center | BDSC: 39171<br>FLYB: FBti0137043 |
| Genetic reagent (*D. melanogaster*) | *He*-Gal4 | Bloomington *Drosophila* Stock Center | BDSC: 8699<br>FLYB: FBst0008699 |
| Plasmids | | | |
| Recombinant DNA plasmid | *pUASz1.0* | (DeLuca and Spradling 2018)<br>Gift of Cooley Lab, Yale | FLYB: FBmc0003175 |
| Recombinant DNA plasmid | pJJH1413 | (Paululat and Heinisch 2012)<br>Gift from the Achim Paululat Lab, Osnabrück University | FLYB: FBmc0003091 |
| Recombinant DNA plasmid | pBluescript KS | Gift of the McKim Lab, Rutgers University | FLYB: FBms0000786 |
| Software | | | |
| Software, algorithm | Imaris | Bitplane | RRID:SCR_007370 |
| Software, algorithm | OriginPro | Origin Lab | RRID:SCR_014212 |

**Table S1. Reagent Table.** Chemicals, fly stocks, plasmids and software used in this study.

| **Table S2: PCR Primers and templates used to generate *pH_TRE_RedStinger_attB*** | | | |
|---|---|---|---|
| PCR fragments used to generate *pH_RedStinger_attB* | | | |
| PCR Product | Forward Primer | Reverse Primer | Plasmid Template |
| attB | TCGACGATGTAGGTCACGGTCTCGA AGC (attB_F) | *TGGCGCGAGCCCCTGATGCTCTTCGTCGACATGC CCGCCGTGACCGTCG (attB_R2)* | *pUASz1.0* (DeLuca and Spradling 2018) |
| 5'_RedStinger | GACGAAGAGCATCAGGGGCTCGCG CC (gyp_hsp70_dsRed_Stinger_2F) | GGACTTGAACTCCACCAGGTAGTGG (RedStinger_ORF_R) | pJJH1413 (Paululat and Heinisch 2012) |
| 3'_RedStinger | CCACTACCTGGTGGAGTTCAAGTCC (RedStinger_ORF_F) | GACGGCGATATTTCTGTGGACAGAGAAGG (ds_RedStinger_R2) | pJJH1413 |
| *miniwhite* | TCCTTCTCTGTCCACAGAAATATCGC CG (Mini-white_F2) | CGCGCCCTAGTTCCAGTGAAATCC (Mini_white_R) | *pUASz1.0* |
| Origin/AmpR | GGATTTCACTGGAACTAGGGCGCGt ttcggggaaatgtgcgcggaacc (Amp_Ori_F) | Tcgagaccgtgacctacatcgtcgaacagaatcaggggataacg cagg (Amp_Ori_R) | pBluescript KS |
| PCR Primers used to amplify TRE sequence | | | |
| TRE_seq_F2 | CAATTGTGCTCGGCAACAGCATGCG | --- | Genomic DNA |
| TRE_R_XhoI | --- | GCCTCTATTTATACTCCGGCGCTCC | Genomic DNA |

**Table S2. PCR Primers and templates used to generate *pH_TRE_RedStinger_attB*.** PCR products, primers and templates used to generate *pH_TRE_RedStinger_attB*. Primers are written in the 5' to 3' direction. Generic Barber lab names are provided for PCR fragments and for primer names found in parenthesis. Genomic DNA was from *TRE_dsRed* transgenic flies.

| Table S3: Substack parameters by glial subtype | | | |
|---|---|---|---|
| **Subtype** | **# Substacks** | **Region** | **Thickness (microns)** |
| **Perineural glia (PG)** | 2 | Anterior central brain region<br>Posterior central brain region | 10-15<br>10-15 |
| **Sub-Perineural glia (SPG)** | 2 | Anterior central brain region<br>Posterior central brain region | 15-20<br>15-20 |
| **Cortex glia (CG)** | 2 | Anterior central brain region<br>Posterior central brain region | 15-20<br>15-20 |
| **Astrocyte-like glia (ALG)** | 1 | Whole central brain | 50-60 |
| **Neuropil ensheathing glia (NEG)** | 1 | Ellipsoid body<br>Internal brain region | 7.5-10 |
| **Tract ensheathing glia (TEG)** | 1 | Ellipsoid body<br>Internal brain region | 7.5-10 |

**Table S3. Substack parameters by glial subtype.** Whole brain images were made into substacks prior to Imaris colocalization analysis which differed depending upon the glia subtype being examined. The number, location, and thickness of the substacks used for different glia subtypes is listed.

| Table S4: Spot XYZ parameters for GFP by glial subtype | | |
|---|---|---|
| **Subtype** | **XY diameter (µM)** | **Z diameter (µm)** |
| **PG** | 5.0 | 10.0 |
| **SPG** | 10.0 | 20.0 |
| **Cortex** | 5.5 | 13.0 |
| **ALG** | 5.0 | 15.0 |
| **NEG** | 5.0 | 10.0 |
| **TEG** | 5.0 | 10.0 |

**Table S4. Spot XYZ parameters for GFP by glial subtype**. To reduce erroneous spot assignment to background or tissue abnormalities, Imaris spot assignment to $GFP^+$ nuclei was limited to parameters that correspond to the expected nuclei size range of the tested glial subtype.

| **Table S5: number of GFP$^+$ glia subtype spots used for colocalization analysis per area analyzed.** | | | | |
|---|---|---|---|---|
| **Subtype** | **Sham/Injury** | **Region** | **# of Regions** | **Mean # of GFP$^+$ nuclei per region** |
| PG | Sham | Anterior | 12 | 230.9 ± 19.5 |
| PG | 1.3 V | Anterior | 3 | 160.3 ± 15.0 |
| PG | 1.6 V | Anterior | 3 | 132.0 ± 19.8 |
| PG | 1.9 V | Anterior | 5 | 173.2 ± 17.6 |
| PG | 2.4 V | Anterior | 10 | 158.7 ± 22.0 |
| PG | Sham | Posterior | 12 | 180.8 ± 7.5 |
| PG | 1.3 V | Posterior | 3 | 144.7 ± 13.9 |
| PG | 1.6 V | Posterior | 3 | 145.0 ± 16.4 |
| PG | 1.9 V | Posterior | 5 | 266.4 ± 27.7 |
| PG | 2.4 V | Posterior | 10 | 156.5 ± 17.7 |
| SPG | Sham | Anterior | 4 | 39.5 ± 6.4 |
| SPG | 1.3 V | Anterior | 4 | 24.8 ± 1.7 |
| SPG | 1.6 V | Anterior | 4 | 34.0 ± 4.1 |
| SPG | 1.9 V | Anterior | 4 | 39.5 ± 3.6 |
| SPG | 2.4 V | Anterior | 4 | 20.8 ± 2.6 |
| SPG | Sham | Posterior | 4 | 35.5 ± 5.7 |
| SPG | 1.3 V | Posterior | 4 | 38.8 ± 4.7 |
| SPG | 1.6 V | Posterior | 4 | 29.0 ± 0.9 |
| SPG | 1.9 V | Posterior | 4 | 37.3 ± 5.1 |
| SPG | 2.4 V | Posterior | 4 | 18.8 ± 3.0 |
| CG | Sham | Whole central brain | 8 | 419.3 ± 75.3 |
| CG | 2.4 V | Whole central brain | 8 | 245.3 ± 57.6 |
| ALG | Sham | Whole central brain | 4 | 644.0 ± 22.5 |
| ALG | 2.4 V | Whole central brain | 4 | 510.8 ± 51.9 |
| NEG | Sham | Ellipsoid body Internal brain region | 8 | 272.8 ± 12.3 |
| NEG | 2.4 V | Ellipsoid body Internal brain region | 9 | 292.1 ± 36.5 |
| TEG | Sham | Ellipsoid body Internal brain region | 4 | 245.3 ± 5.0 |
| TEG | 2.4 V | Ellipsoid body Internal brain region | 4 | 212.5 ± 10.1 |

**Table S5. Number of GFP$^+$ glia subtype spots used for colocalization analysis per area analyzed.** Number of GFP$^+$ glia subtype spots tested for colocalization with RedStinger differed depending upon which glia subtype was being tested.

**Figure S1.**

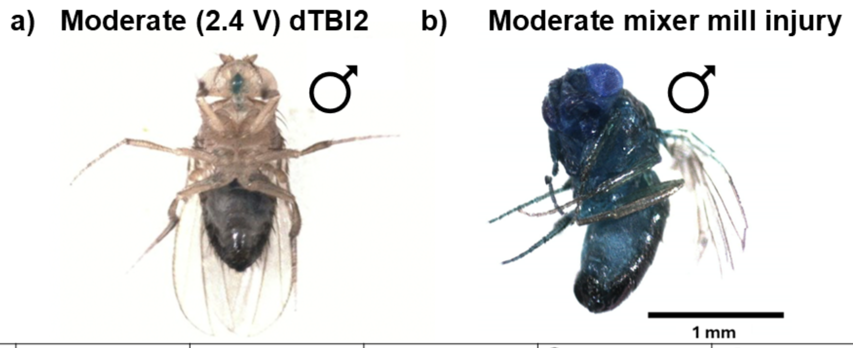


c)

| | Flies per injury | Flies alive post-injury | # Smurf post-injury | ♀ Smurfs | ♂ Smurfs | % Smurfs |
|---|---|---|---|---|---|---|
| Dyed food uninjured | 18 | 0 | 0 | 0 | 0 | 0 |
| Dyed food injured | 30 | 14 | 4 | 1 | 3 | 28.5% |

**Figure S1. dTBI2-induced mortality is not due to gut barrier leakage.** Flies were fed a less nutritive dyed diet beginning 2 days prior and ending 24 h post-injury. a) Representative male fly subjected to moderate 2.4 V injury using dTBI2, showing no smurfing as dye is consolidated in the gut. b) Representative male fly subjected to moderate TBI using mixer mill at 30 seconds and 100 rpm, showing smurfing as gut barrier integrity is disrupted leading to dye leaking throughout the body. c) Table of experimental information and percent of flies with gut barrier disruption post-injury using mixer mill only.

**Figure S2.**

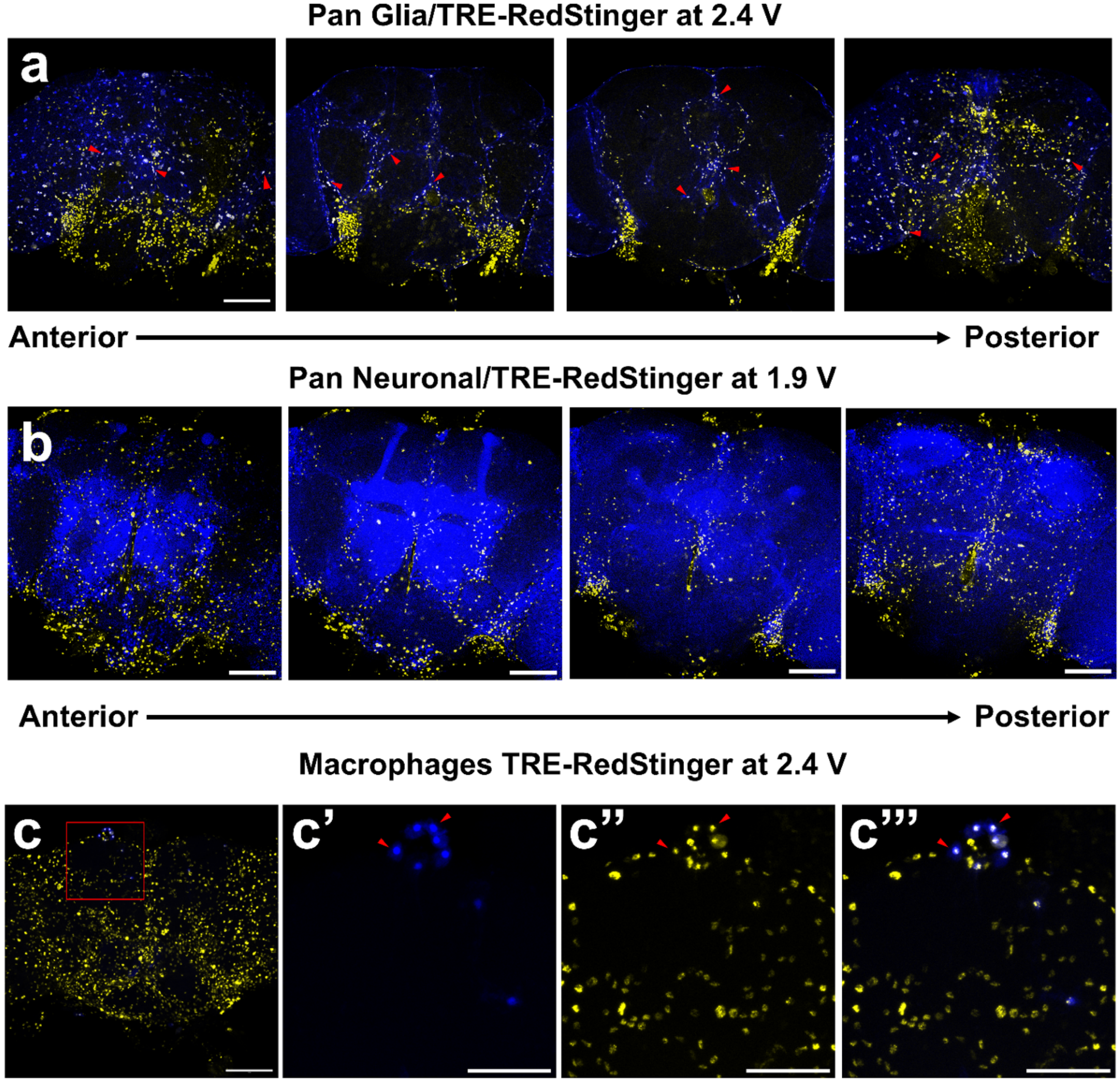


**Figure S2. Non-glial and non-neuronal cells show AP-1 activation after TBI.** For all image panels, brains were dissected 24 h post injury, GFP (blue) and RedStinger (yellow), colocalization of GFP and RedStinger (white). a) Moderate 2.4 V injury, *TRE_RedStinger/*+; *repo-Gal4*$^{M1B}$/*UAS-GFP.nls.* b) Mild 1.9V injury, *TRE_RedStinger/*+; *nSyb*-Gal4/*UAS-GFP.nls*, (a, b) Single serial confocal sections of a single brain arranged from the antennal anterior side (left) to the posterior side (right) of the brain. (c, c', c'', c''') *TRE_RedStinger/*+; *He-Gal4*/*UAS-GFP.nls*, moderate 2.4 V injury, magnification of area marked by the red square in panel (c) and displayed as separate (c') GFP, (c'') RedStinger and (c''') merged panels. Red arrows indicate examples of colocalized nuclei in all panels. (a,b,c) Scale bar = 100 microns (c',c'',c''') Scale bar = 50 microns.

**Figure S3.**

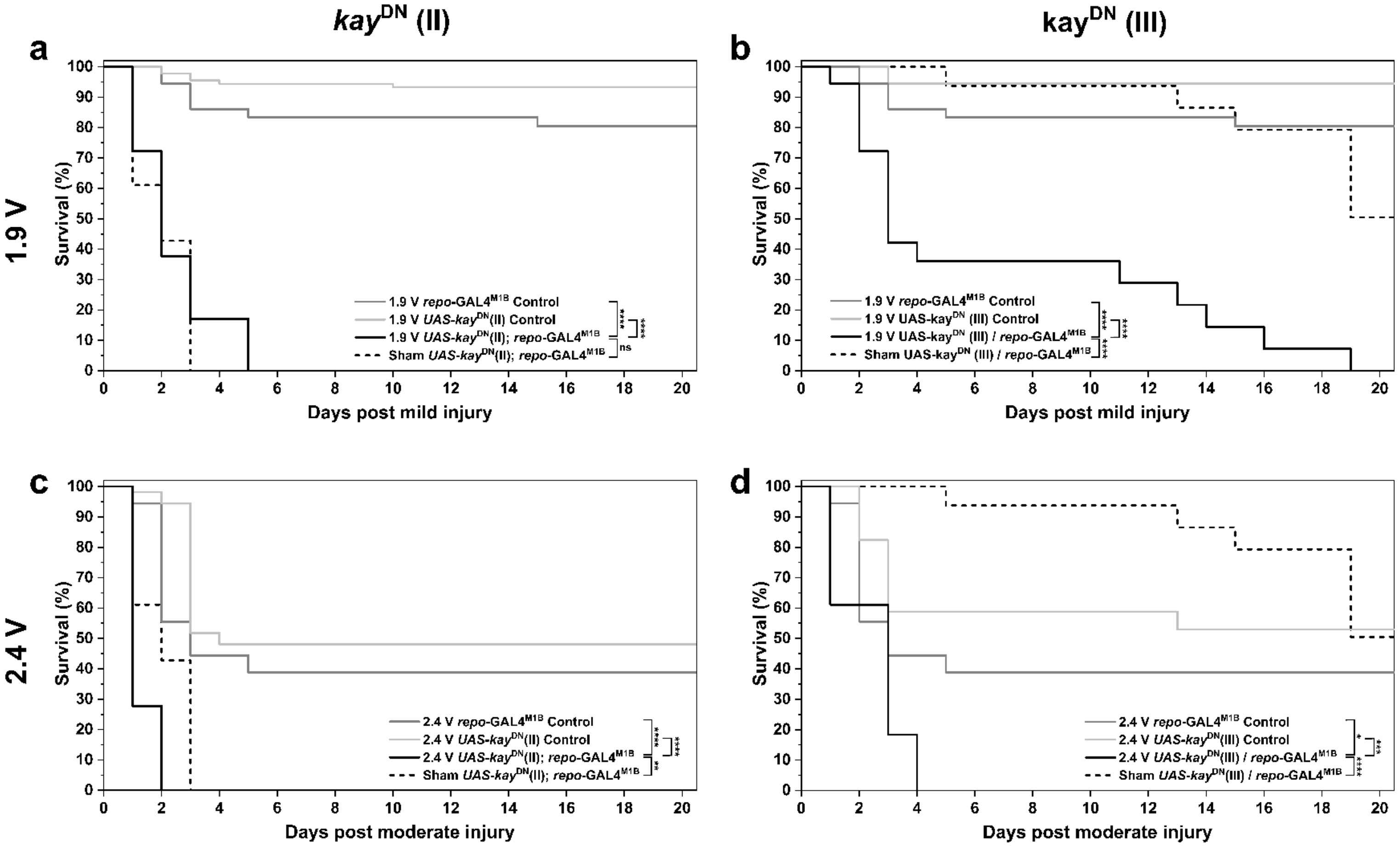


**Figure S3. Pan-glial expression of UAS-*kay*$^{DN}$ shows variable mortality phenotypes depending upon chromosomal location of transgene.** Twenty-one day survival experiments examining effects of injury on flies expressing either *UAS-kay*$^{DN}$ (II) or *UAS-kay*$^{DN}$ (III) by the pan-glia *repo-Gal4*$^{M1B}$ driver. (a, b) mild 1.9 V injury (c, d) moderate 2.4 V injury. (a, b, c, d) *repo-Gal4*$^{M1B}$/+ control (dark gray), (a, c) *UAS-kay*$^{DN}$ (II) /+ control (light gray), (b, d) *UAS-kay*$^{DN}$ (III) /+ control (light gray), (a, c) *UAS-kay*$^{DN}$(II) /+; *repo-Gal4*$^{M1B}$ /+ experimental (black), (b, d) *repo-Gal4*$^{M1B}$ /*UAS-kay*$^{DN}$ (III) experimental (black), (a, b, c, d) Experimental Shams (dashed black). N = 18 flies per condition. For all survival experiments, Sham flies were collared, left in the collar several minutes then removed with forceps. Log-rank pairwise comparison was used to determine statistical differences between injured groups for survival assays. Brackets in legend and graphs designate the statistical significance between data sets, ns = not significant, * $p<0.05$, ** $p<0.01$, *** $p<0.001$, **** $p<0.0001$.
